# Identification of Antituberculars with Favorable Potency and Pharmacokinetics through Structure-Based and Ligand-Based Modeling

**DOI:** 10.1101/2025.02.03.636334

**Authors:** Vedang Warapande, Fanwang Meng, Alexandra Bozan, David E. Graff, Jenna C. Fromer, Khadija Mughal, Faheem K. Mohideen, Shivangi, Sindhuja Paruchuri, Melanie L. Johnston, Pankaj Sharma, Timothy R. Crea, Reshma S. Rudraraju, Amir George, Camilla Folvar, Andrew M. Nelson, Matthew B. Neiditch, Matthew D. Zimmerman, Connor W. Coley, Joel S. Freundlich

## Abstract

Drug discovery is inherently challenged by a multiple criteria decision making problem. The arduous path from hit discovery through lead optimization and preclinical candidate selection necessitates the evolution of a plethora of molecular properties. In this study, we focus on the hit discovery phase while beginning to address multiple criteria critical to the development of novel therapeutics to treat *Mycobacterium tuberculosis* infection. We develop a hybrid structure- and ligand-based pipeline for nominating diverse inhibitors targeting the β-ketoacyl synthase KasA by employing a Bayesian optimization-guided docking method and an ensemble model for compound nominations based on machine learning models for *in vitro* antibacterial efficacy, as characterized by minimum inhibitory concentration (MIC), and mouse pharmacokinetic (PK) plasma exposure. The application of our pipeline to the Enamine HTS library of 2.1M molecules resulted in the selection of 93 compounds, the experimental validation of which revealed exceptional PK (41%) and MIC (19%) success rates. Twelve compounds meet hit-like criteria in terms of MIC and PK profile and represent promising seeds for future drug discovery programs.

## INTRODUCTION

Tuberculosis (TB), caused by *Mycobacterium tuberculosis*, poses a major threat to global health, with 10.8 million cases leading to 1.3 million deaths in 2023 alone^1^. The incidence rate of TB exhibits a significant correlation with socioeconomic conditions^2^. The treatment of TB commonly involves the oral administration of multiple drugs for a period of approximately 6 months, challenging patient adherence. Therapeutic interventions are further complicated by varying degrees of drug resistance^3^, drug adverse effects^4^, and the potential for drug-drug interactions^5^. Antitubercular drugs with improved efficacy, lack of cross-resistance with existing therapies, and the potential to shorten the duration of a regimen may offer improved treatments for this multi-decade pandemic.

The cell wall of *M. tuberculosis* plays a crucial role in its virulence and represents an array of validated drug targets^6^. Its mycolic acid layer is comprised of long chains of α-alkyl, β-hydroxy fatty acids which are synthesized and elongated in part by the type Ⅱ fatty acid synthase (FAS-Ⅱ) pathway. The approved drug isoniazid (INH) inhibits the FAS-Ⅱ enoyl-acyl carrier protein reductase InhA^7, 8^. Additional efforts have identified the β-ketoacyl synthase KasA, another enzyme in the FAS-Ⅱ pathway, as an essential and vulnerable protein target in *M. tuberculosis*^9,10^. We have recently disclosed a preclinical candidate for the treatment of TB, JSF-3285, that inhibits KasA and achieves significant *in vitro* and *in vivo* potency against *M. tuberculosis*^11^. Requisite to the translation of a KasA inhibitor to clinical trials, JSF-3285 is currently in preclinical development studies while we are tasked with the pursuit of second-generation KasA inhibitors. The identification of different chemical classes, or chemotypes, will be key to the attainment of next-generation molecules that maintain, or improve upon, the promising efficacy and pharmacokinetic profiles of JSF-3285.

Drug discovery can be understood as a multi-parameter optimization problem^12–14^, where *in vitro* efficacy and rodent PK are two important objectives. In particular, consideration of compound PK profile metrics, such as compound exposure with oral dosing, at the earlier stages of drug discovery can reduce the risk of failure during the later phases^15–17^. Herein we describe a computational pipeline (Figure 1) that merges approaches in structure-based design^18, 19^ and machine learning models for molecular property predictions^20, 21^ to identify diverse small molecule growth inhibitors of *M. tuberculosis*. First, the Enamine HTS (2.1M) collection (2020 release; www.enamine.net) was screened with an active learning strategy^22^ to identify compounds predicted to bind to KasA based on docking score. The prioritized candidates were filtered with ligand-based binary classification machine learning models to predict *in vitro* growth inhibition of *M. tuberculosis* (as quantified by minimum inhibitory concentration (MIC)), representing the lowest compound concentration to achieve consequential growth inhibition of the bacterium in culture (at the 90% level), and mouse plasma exposure (as quantified by AUC_0-5h_, the area under the curve for 0 – 5 h) post a single oral (po) dose. Experimental testing of 93 candidates revealed 38 as meeting the criterion of PK AUC_0-5h_ ≥ 1,000 h*ng/mL and 18 meeting the criterion of MIC ≤ 12 µM. Of particular interest are 12 compounds that satisfy both objectives and have the potential to seed *M. tuberculosis* KasA drug discovery efforts.

**Figure 1.**
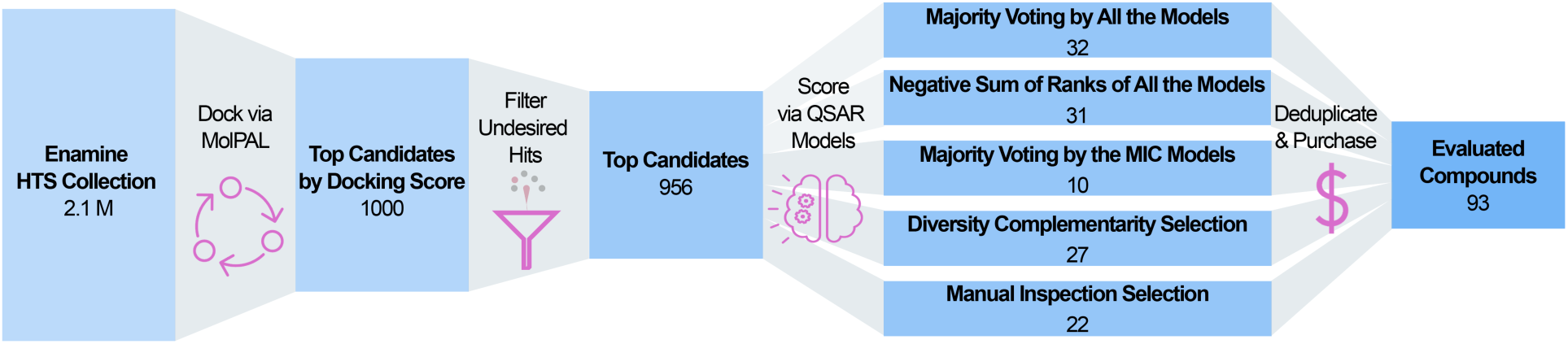
Computational workflow for prioritizing hit compounds for experimental validation. Beginning with the 2.1M Enamine HTS Collection compounds, an active learning-based docking tool MolPAL provided the top 1,000 candidates prioritized by docking score. Compounds with substructure alerts were removed, leaving 956 candidates. Five different compound nomination strategies were utilized to identify 96 compounds for testing after deduplication. Of these, 93 candidates were successfully sourced commercially and experimentally profiled.

## RESULTS

### Model-guided virtual screening identifies putative KasA binders

Compounds were first prioritized for experimental assessment through a structure-based virtual screen to identify putative inhibitors of KasA. Limitations in computational resources, combined with the hypothesis that “bigger is better” in structure-based virtual screening^23–26^, motivated the use of an active learning approach to efficiently explore the Enamine HTS Collection compound set containing roughly 2.1M molecules. Specifically, we applied MolPAL (Molecular Pool-based Active Learning^22^) to iteratively explore the library by docking batches of ∼8.2k compounds. In each iteration, a graph neural network (GNN) was trained on the acquired dataset of compounds and associated Vina docking, starting from a randomly acquired initialization set. The library was then rank-ordered by predicted scores and the molecules with the lowest predicted scores were then selected for docking. This approach ultimately resulted in docking 51k compounds, with the top-1,000 compounds all possessing Vina docking scores ≤ −10.9 kcal/mol. These top-scoring compounds were filtered to remove those with substructures associated with DNA intercalation^28^ and potential mutagenicity/genotoxicity^29, 30^, yielding a final collection of 956 (Supplementary File S1).

### Machine learning models provide complementary criteria for downselection

To prioritize compounds from the 956 identified by model-guided virtual screening, we trained machine learning models to predict mouse PK exposure and *in vitro* growth inhibition of *M. tuberculosis*. Classifications models were used to predict PK AUC_0-5h_ from our in-house PK dataset^31^ consisting of 190 unique compounds, classifying all compounds with mouse plasma AUC_0-5h_ ≥ 1,000 h*ng/mL as PK active. We identified three publicly available datasets for *M. tuberculosis* growth inhibition: TAACF-Chembridge^32^, TAACF-Kinase^33^, and MLSMR^34^. Both TAACF datasets contain single-point percent growth inhibition for compounds at 10 µg/mL concentration and were therefore merged into a single, non-redundant dataset (“MIC-TAACF”) of 120K compounds. The MLSMR dataset (“MIC-MLSMR”) contains single-point growth inhibition measurements at a 10 μM compound concentration for 214K compounds. We trained separate classification models on the MIC-TAACF and MIC-MLSMR datasets, using a threshold of ≥90% inhibition for both antitubercular activity models.

We pursued several tree-based and deep learning architectures, including random forest (RF), XGBoost, LightGBM, CatBoost, ChemBERTa-v2^35^, and a directed message passing neural network (D-MPNN)^21, 36^. Tree-based models used the Molecular Operating Environment (MOE version 2022.02) 2D descriptors and Morgan fingerprints^37^ as features. For each dataset, the best model architectures and training hyperparameters were determined using hyperparameter tuning with various objective functions such as binary cross entropy (BCE) Loss, area under the Receiver Operating Characteristic curve (AUROC), average precision (AP), and the maximum F1 score across all thresholds (max. F1 score). Further details related to dataset curation, model training, and hyperparameter tuning may be found in the Materials and Methods section. Models were evaluated according to performance on a held-out test set in terms of their test AUROC, AP, and max. F1 scores (Table 1). We forwent a formal model comparison using statistical significance testing^38–41^, as these surrogate models were intended for use in the subsequent prioritization of the 956 docking-derived candidates and not for drawing generalizable conclusions about model performance. Top performing models were selected based on their mean performance, as the model with the best mean performance is unlikely to be significantly worse than another model. In the case of the PK dataset, the random forest and gradient-boosted tree models generally exhibited higher performance metrics than the deep learning models other than the D-MPNN models. These numerical trends were also observed for the models of the two MIC datasets. We chose specific model instances (Tables S1 and S2) of these best performing algorithms based on their test AUROC and AP for downstream candidate selection.

**Table 1.** Performance of select models on the PK, MIC-TAACF, and MIC-MLSMR datasets. The metrics reported for model comparisons (AUROC, AP, and max F1 score) were calculated on the test split of the dataset. An underlined entry indicates the model with the largest value for the pertinent metric.

| Model | Objective<br>Function | AUROC | Avg.<br>Precision<br>(AP) | Max F1 Score |
| --- | --- | --- | --- | --- |
| PK (190) |  |  |  |  |
| <u>Random Forest</u> | AUROC | <u>91.4</u> | <u>87.7</u> | <u>90.0</u> |
| XGBoost | AUROC | 90.2 | 82.9 | 88.9 |
| LightGBM | AUROC | 88.0 | 83.3 | 84.4 |
| CatBoost | AUROC | 89.2 | 86.8 | 85.0 |
| ChemBERTa-V2 | BCE Loss | 78.7 | 80.1 | 76.9 |
| D-MPNN | BCE Loss | 85.0 | 86.1 | 78.3 |
| MIC-TAACF<br>(120K) |  |  |  |  |
| <u>Random Forest</u> | F1 Score | 89.0 | <u>34.2</u> | 38.6 |
| XGBoost | AP | 87.8 | 33.9 | <u>40.8</u> |
| LightGBM | AP | 87.2 | 32.2 | 40.5 |
| CatBoost | F1 Score | 85.5 | 31.5 | 38.5 |
| ChemBERTa-V2 | BCE Loss | 85.2 | 31.9 | 38.2 |
| D-MPNN | BCE Loss | <u>89.7</u> | 29.5 | 38.4 |
| MIC-MLSMR<br>(214K) |  |  |  |  |
| <u>Random Forest</u> | F1 Score | 87.2 | <u>29.8</u> | 36.4 |
| XGBoost | AP | 86.6 | 27.3 | 32.7 |
| LightGBM | AP | 84.9 | 21.7 | 28.3 |
| CatBoost | AP | 84.0 | 26.2 | 32.8 |
| ChemBERTa-V2 | BCE Loss | 56.1 | 2.5 | 0.0 |
| D-MPNN | BCE Loss | <u>88.6</u> | 29.1 | <u>36.6</u> |

### Ligand-based scoring and pose inspection affords a prioritized set of 96 candidates for experimental testing

The 956 docking-derived candidates were further prioritized using multiparameter scores that considered our model predictions of PK plasma exposure and *in vitro* antitubercular activity. These model instances (Tables S1 and S2) provided labels for the 956 compounds indicating predicted PK and MIC activity. We defined three multiparameter scores: (1) Majority Voting by All the Models, (2) Negative Sum of Ranks of All the Models, and (3) Majority Voting by the MIC models only, respectively (Figure 1). Further details as to these nomination protocols may be found in Materials and Methods and in Supporting Information. We selected 31, 31, and 10 compounds with these three methods, respectively. Once duplicate chemical structures were removed, we arrived at a set of 47 candidates. To ensure that the chemical space of the putative KasA binders was sufficiently represented, the 956 candidates were clustered based on Morgan fingerprints (1024 bits, radius = 2) into 20 clusters, nine of which were not represented by previous selections. Using what we refer to as a Diversity Complementarity approach, we selected 3 compounds from each unrepresented cluster, leading to an additional 27 candidates. An additional 22 compounds were selected manually by inspection of their docked poses with KasA. Termed Manual Inspection in Figure 1, this nomination strategy was guided by the reported protein-ligand interactions of JSF-3285^11^ to include: (1) proposed hydrogen bonding with Glu199 with a maximum distance of 3.6 Å as well as other hydrogen bonding criteria from PyMOL; (2) selecting for a bound pose with a reasonable, visual alignment with JSF-3285; (3) proposed formation of additional hydrogen bonds to one or more of the following residues (Glu120, Pro201, Glu203) (with the same distance criterion as for (1)); (4) visually identified (centroid to centroid distance ∼ 3.5 – 4.5 Å ^42^) face-to-face pi-pi stacking interactions with one or more of the following residues (Phe210, Phe239, Tyr82); (5) a preference for compounds with a lower molecular weight. In total, 96 compounds were nominated with 93 available from the commercial vendor for purchase and subsequent experimental testing (Supplementary File S1).

### Experimental validation shows enrichment for antitubercular activity and favorable PK profile

93 of the 96 selected compounds were sourced from Enamine Ltd. (www.enamine.net) and evaluated in the mouse snapshot PK model^43, 44^ with female CD-1 mice. An impressive 41% (38/93 candidates) met the acceptable criterion of AUC_0-5h_ ≥ 1,000 h*ng/mL (Figure 2, Supplementary File S1). Eleven compounds were not profiled due to insufficient stability in mouse plasma (plasma half-life t_1/2_ ≤ 30 min) that precluded the construction of a calibration curve, and one additional compound was not sufficiently soluble to be assayed for mouse PK. The acquired compounds were also evaluated for their growth inhibition of the *M. tuberculosis* H37Rv strain (Figure 2, Supplementary File S1). 18/93 (19%) compounds exhibited an acceptable MIC ≤ 12 μM. INH was used as a positive control (MIC = 0.078 μM). Two candidates were of insufficient solubility to enable an MIC determination. It should be noted that the MIC-TAACF models utilized data with µg/mL units while the MIC-MLSMR model data had μM units. Thus, we chose to utilize the more lenient 12 μM cutoff given that drug-like small molecules typically have a molecular weight ≤ 500 g/mol. Finally, the intersection of candidates with acceptable PK AUC_0-5h_ and MIC values afforded 12 hit compounds (12/93 = 13% hit rate) deemed worthy of further study.

**Figure 2.**
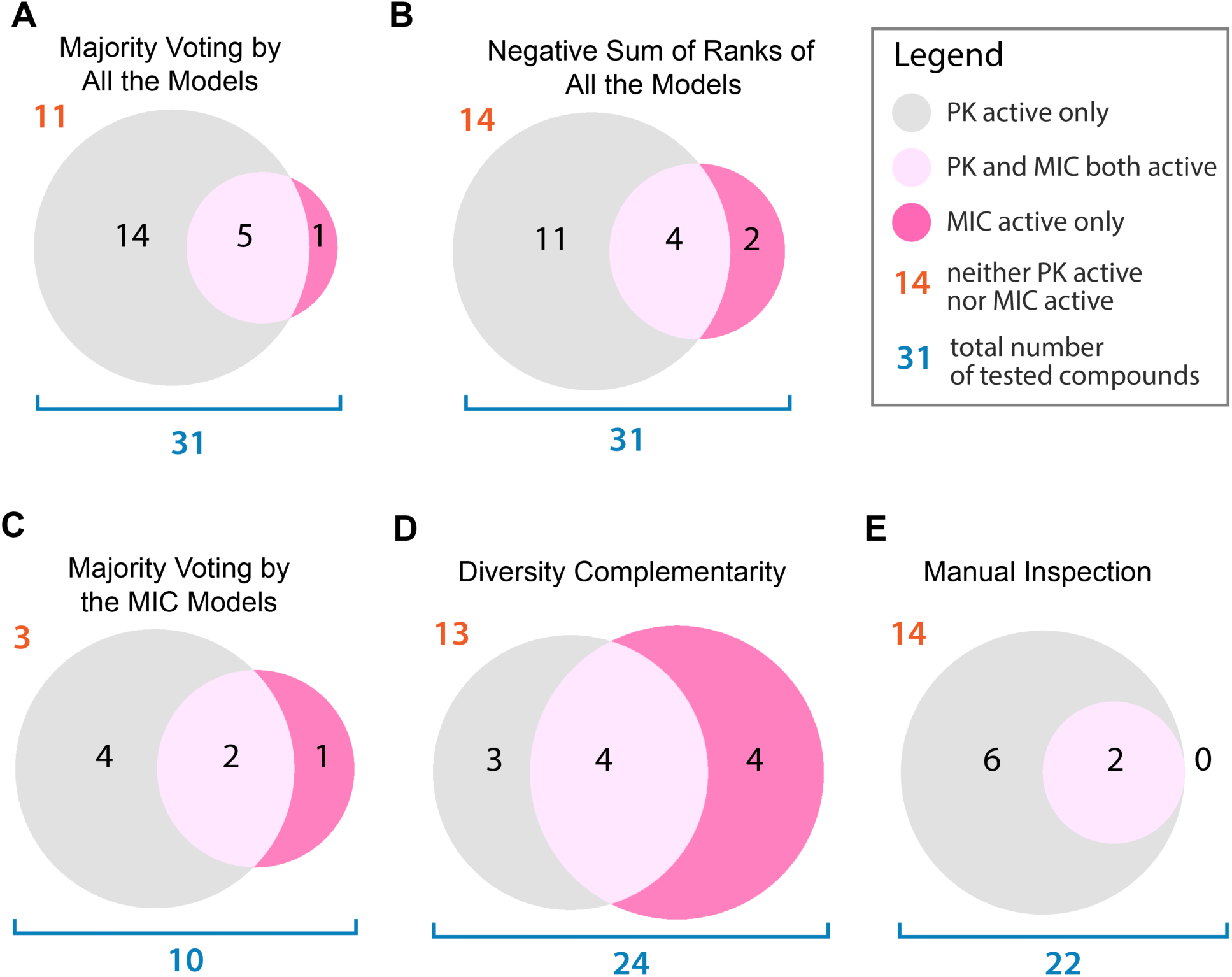
Venn diagrams of the PK and MIC active compounds for each compound nomination strategy, (A)-(E). Bioactive molecules are defined with MIC ≤ 12 μM and/or PK AUC_0-5h_ ≥ 1,000 h*ng/mL. The legend defines the meaning of each panel for the example of strategy B. In some cases, candidates were chosen by more than one model (Supplementary File S1). While 27 compounds were selected from the Diversity Complementarity method, only 24 compounds were sourced as 3 compounds were unavailable due to excessive cost and/or changes in the Enamine HTS Collection between the time of download and the time of purchase.

Visual inspection of the twelve hits facilitates their clustering (Figure 3). The first cluster features 3 compounds (Z31753778, Z318114792, Z979305278) which share the (3-trifluoromethyl)benzene group on one side, joined through a combination of amide/sulfonamide linkages to a terminal aromatic moiety. The 4 compounds in the second cluster (Z29466477, Z99494139, Z99494149, Z212116604) share a 2-arylquinoline 4-amide substructure with Z99494139 and Z99494149 differing only by the 4-substituent of the 2-aryl moiety. The third cluster contains two other 2,4-disubstituted quinolines (Z1143441182, Z56802551). The remaining compounds are the 2,3,4-substituted quinoline Z1702969146, quinazoline Z1033301210, and benzimidazol-2-one Z126939482. To the best of our knowledge, none of the twelve hits has previously been disclosed as demonstrating antitubercular growth inhibitory activity. We do note that other 2-substituted quinoline-6-amides have been reported to be growth inhibitors of *in vitro* cultured *M. tuberculosis*^45^.

**Figure 3.**
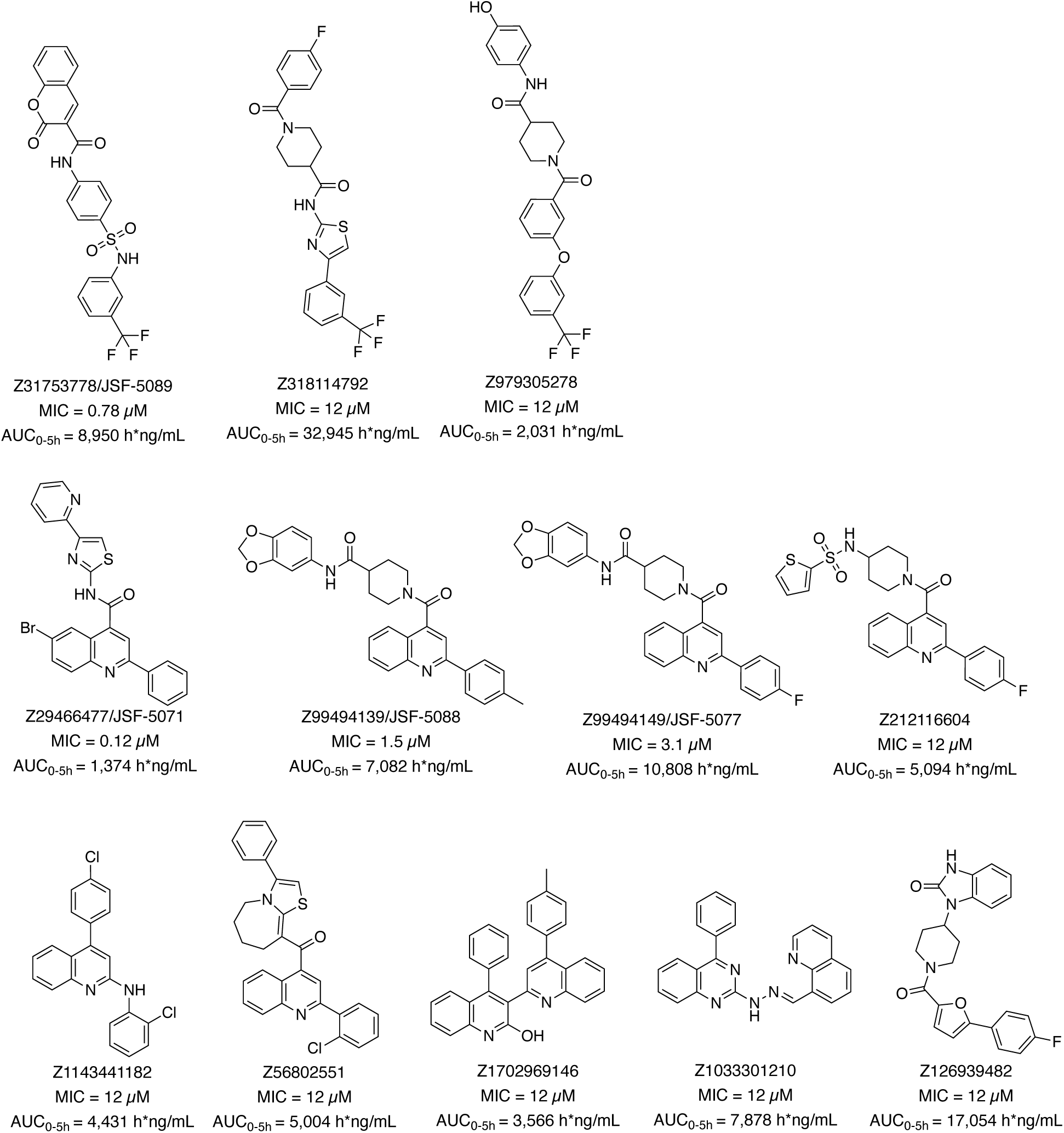
The chemical structures of the 12 hit compounds with their identifiers, MIC, and AUC_0-5h_ values.

### KasA target engagement and *in vivo* profiling of the hit compounds

We assessed the propensity of these 12 hits to inhibit the catalytic function of purified KasA. Inhibition was determined via a functional assay coupling KasA activity to the next enzyme in the FAS-Ⅱ pathway, the NADPH-dependent β-ketoacyl-AcpM reductase MabA^46–49^. Compounds were initially assessed for percent inhibition at a single concentration of 25 µM (Figure 4; Table S3). 3/12 compounds (Z979305278, Z1033301210, and Z1702969146) demonstrated modest (≥25%) functional inhibition of purified KasA, as compared to JSF-3285 as a positive control with 92.6 ± 3.5% inhibition. Counterscreening versus MabA showed that 11/12 compounds at a 25 µM concentration exhibited negligible (≤10%) inhibition of 500 nM MabA (Figure S1, Table S4). As negative controls, JSF-3285 and DMSO each exhibited zero percent inhibition of MabA under the assay conditions. The one compound (Z1033301210) with modest inhibition (31.6%) of MabA was further characterized with an IC_50_ > 100 µM versus MabA. This negligible inhibition of MabA is at an enzyme concentration 10% of that in the coupled enzyme assay.

**Figure 4.**
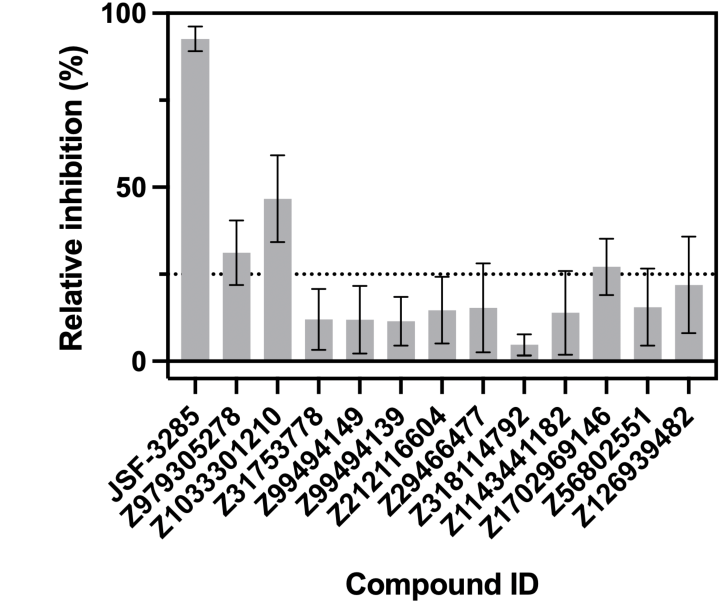
KasA functional inhibition for the 12 hit compounds (at 25 µM). Reported values represent the mean ± standard deviation from n = 6 independent trials. Relative inhibition is calculated compared to a no inhibitor control.

### Synthesis of select hits and their further biological profiling

Amongst the 12 hits, the four most potent growth inhibitors of *in vitro* cultured *M. tuberculosis* were synthesized (Scheme 1). Z29466477 was synthesized as JSF-5071, relying on the Pfitzinger reaction^50, 51^ of commercially available 5-bromoindoline-2,3-dione with acetophenone followed by HATU-mediated coupling^52^ with 4-(pyridin-2-yl)thiazol-2-amine. The structurally related Z99494149 and Z99494139 were prepared as JSF-5077 and JSF-5088, respectively. Their shared N-(benzodioxol-5-yl)piperidine-4-carboxamide moiety was prepared via the HATU-mediated coupling of commercially available benzo[d][1,3]dioxol-5-amine and 1- (*tert*-butoxycarbonyl)piperidine-4-carboxylic acid, followed by Boc-deprotection with trifluoroacetic acid. The resulting amine was coupled with the appropriate 2-substituted quinoline-4-carboxylic acid (4-fluorophenyl for JSF-5077 and 4-methylphenyl for JSF-5088), prepared by Suzuki-Miyaura cross-coupling^53^ of commercial methyl 2-chloroquinoline-4-carboxylate with the requisite 4-substituted phenylboronic acid followed by saponification of the methyl ester. Z31753778 was synthesized as JSF-5089 through the following route: reaction of 4-nitrobenzenesulfonyl chloride with 3-(trifluoromethyl)aniline, reduction of the nitroarene to the aniline with Zn/NH_4_Cl^54^, and HATU-mediated coupling with commercial 2-oxo-2H-chromene-3-carboxylic acid.

**Scheme 1.**
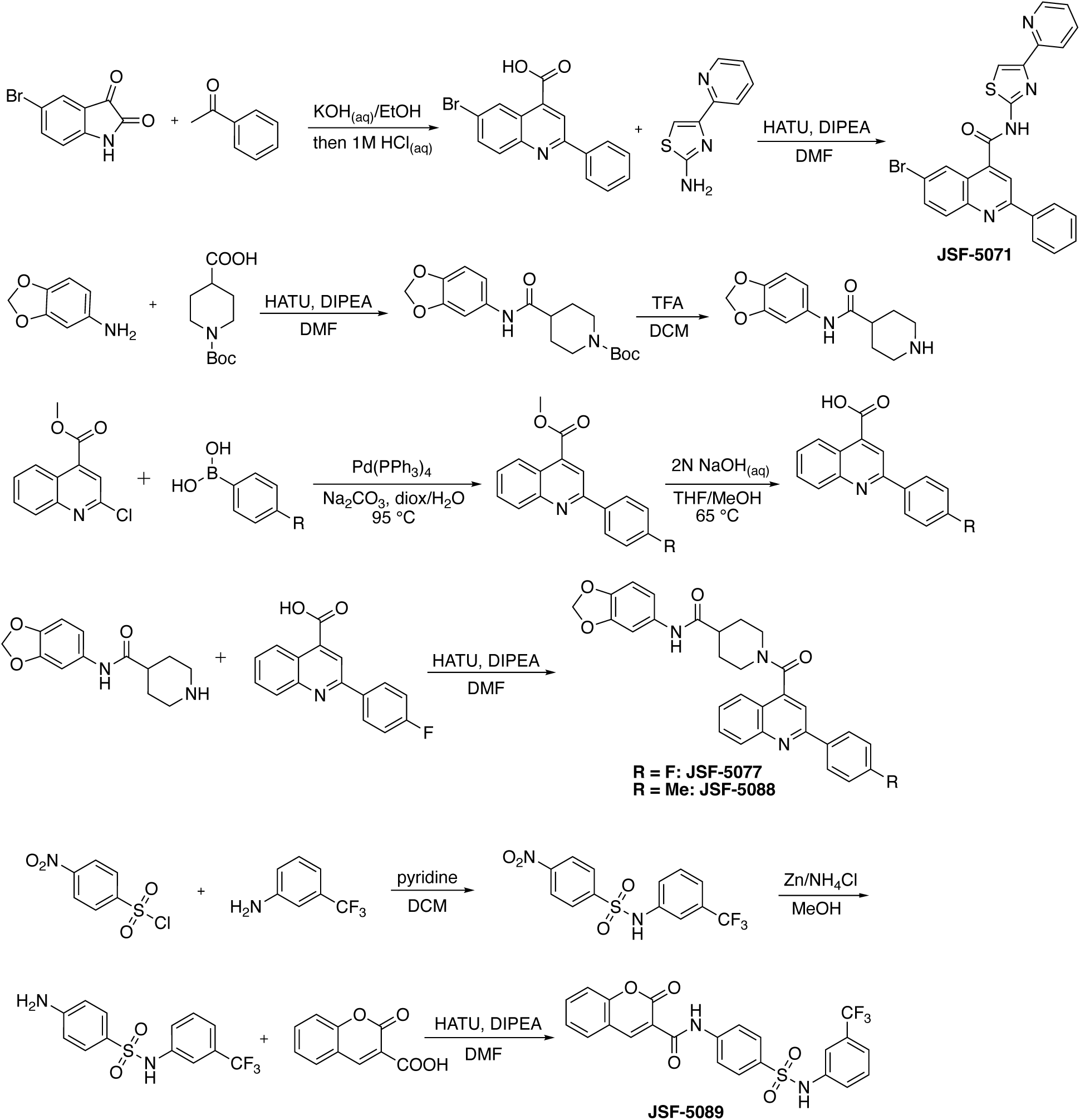
Synthetic routes for the preparation of select hit compounds.

These four synthesized hits exhibited similar (within ±1 dilution) *M. tuberculosis* MIC values as observed with the commercially sourced compounds. Given that JSF-5071 exhibited the most potent MIC of 0.12 µM, we proceeded with its further profiling. We note that JSF-5071 displayed a CC_50_ = 0.62 – 0.78 µM versus *in vitro* cultured Vero cells (ATCC # CCL-81) where CC_50_ is the minimum compound concentration to afford 50% growth inhibition of this model mammalian cell line. Thus, JSF-5071 has a selectivity index equal to its CC_50_/MIC = 5.2 – 6.5 that is moderate yet close to the desired hit value of 10 ^55^. It exhibited very modest functional inhibition of KasA (12.7 ± 4.8% at 25 µM compound; Table S3) but demonstrated a KasA K_d_ of 4.78 ± 1.3 µM by microscale thermophoresis (MST; Figure 5) as compared to JSF-3285 (K_d_ = 70.7 ± 23.5 nM)^56^. JSF-5071 was assessed for its 24 h PK profile post po and intravenous (iv) dosing in mice (Table S5 and Figure S2). Its oral bioavailability (%F) was 18.5 with a solution formulation of 1:9 DMSO:20% Solutol HS15 (%F = 10.7 with a suspension formulation of 5% DMA/95% CMC/Tween), which is below our typical goal value of 30 ^57^. Dosing above 25 mg/kg was limited by solubility in non-toxic dosing vehicles to enable solution formulation dosing. Thus, a 25 mg/kg once-daily (qd) po dose was assessed in a 4 day tolerability study with a solution formulation using 1:9 DMSO:20% Solutol HS15 (Figure S3). JSF-5071 was well tolerated without weight loss or visible signs of stress in the mice. The day 4 AUC_0-24h_ at steady state was slightly greater than twofold higher than the day 1 AUC_0-24h_ (24 h*µg/mL vs. 10 h*µg/mL). JSF-5071 was studied in a BALB/c mouse model of acute *M. tuberculosis* infection using this solution formulation via low-dose aerosol exposure to further query its efficacy during the log-growth phase of infection^58, 59^. Seven days post-infection with the *M. tuberculosis* Erdman strain, the mice were dosed with 25 mg/kg qd po JSF-5071 for 12 days. The front-line drug rifampicin (10 mg/kg twice-daily (bid) po) and vehicle only were the positive and negative controls, respectively. JSF-5071 did not demonstrate a statistically significant reduction in mouse lung colony-forming units (CFUs) at 21 days post-infection as compared to vehicle treated mice (Figure S4).

**Figure 5.**
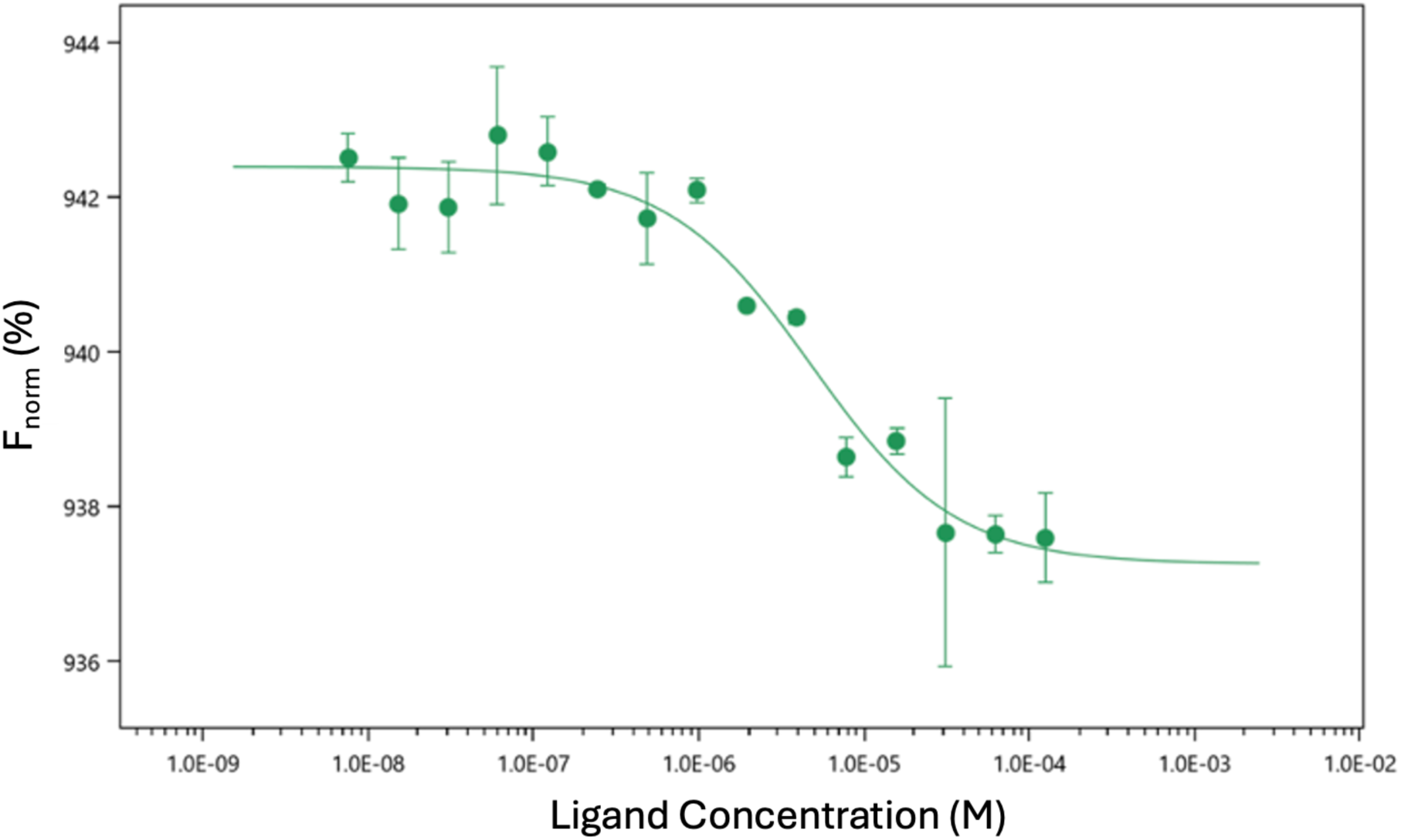
MST quantification of JSF-5071 binding to KasA. The data for JSF-5071 is presented as the mean ± standard deviation of two independent assays.

## DISCUSSION AND CONCLUSIONS

An often employed workflow in computationally focused hit discovery is to perform virtual screening, e.g., docking, experimentally test the top candidates, and then optimize the hits to leads through a focus on one or more properties, including *in vitro* potency, *in vitro* Absorption-Distribution-Metabolism-Toxicity (ADMET) metrics, and rodent PK profile, before entertaining assessment in a first-line rodent model of efficacy^60, 61^. A common challenge is that the computationally identified hit compounds often display certain structural motifs requisite for *in vitro* potency but detrimental to ADMET and/or PK profiles. Drug discovery and development inherently are represented by a multiple criteria decision making problem^62–64^. The array of molecular properties and profiles that must be evolved to be within acceptable limits rises considerably as a program traverses from hit to lead to preclinical candidate. *In vitro* potency (including on-target engagement as quantified by binding affinity and inhibition of the protein’s native function), *in vitro* ADMET, *in vivo* PK, and *in vivo* safety profiles are amongst the most notable criteria. In this work, we disclose our efforts to approach hit discovery while laying the groundwork for the above-mentioned later optimizations.

We have demonstrated a computational pipeline to identify novel antitubercular hit compounds with acceptable values for *in vitro* growth inhibition of *M. tuberculosis* (MIC) and mouse snapshot PK profiles (plasma AUC_0-5h_) and a heightened probability of engaging KasA. Our approach integrates structure- and ligand-based drug discovery methods, beginning with an active learning approach to docking to the JSF-3285 binding pocket, the acyl binding site^11, 18^ of KasA. The use of model-guided docking^22, 23^ enables the consideration of a large initial chemical space, restricted to commercially available molecules to facilitate sourcing and experimental validation of compounds passing subsequent filtration and prioritization stages. After the docking step, the workflow moved to the construction of machine learning models for molecular property predictions with which to filter the hit list. For both the PK and MIC datasets, the random forest, gradient-boosted tree, and D-MPNN models were amongst the top performing.

Five different compound nomination strategies based on combinations of ML models for MIC and PK predictions and structural diversity were implemented to arrive at the 93 selected compounds for biological assay. Prioritization based on three distinct multi-parameter scores (Majority Voting by All the Models, Negative Sum of Ranks of All the Models, Majority Voting by the MIC Models) afforded excellent success rates in predicting MIC (19 – 30%) and PK (48 – 61%) active compounds. The Diversity Complementarity method identified 8/24 (33%) and 7/24 (29%) MIC and PK active compounds, respectively. In addition, this complementary strategy identified 4 out of 12 hit compounds (Z31753778, Z99494139, Z99494149, and Z979305278) that were not selected by other nomination methods, supporting the importance of covering broad chemical spaces and not over-relying on structure-property relationship models expected to exhibit a modest false negative rate. Moreover, the hits Z126939482 and Z212116604 were only selected by manual inspection of docked poses. However, manual selection of compounds without consideration of ML property-based models had the lowest success rate for MIC prediction (2/22 = 9%).

One would expect that the use of docking alone would predict, at best, target binding in a biophysical assay. MIC is a function also of the degree of inhibition of the target’s native function that is tied to the growth of the bacterium. Furthermore, docking does not account for the drug accumulation barrier, which is significant in *M. tuberculosis* due to its complex and extremely hydrophobic cell wall composition^65, 66^. However, we do note that 8/22 or 36% of the Manual Inspection hits were PK actives. This result may be due to luck alone or perhaps reflect on the nature of the KasA binding pocket and the inadvertent correlation of favorable PK and docking scores. Finally, in consideration of their ability to predict the 12 hits, the rank order of the nomination methods was: Majority Voting by All the Models (5), Negative Sum of Ranks of All the Models (4 correct), Diversity Complementarity (4), Majority Voting by the MIC Models (2), and Manual Inspection of Docked Modes (2).

The twelve hit compounds may serve as starting points for further hit expansion and *in vitro* ADMET property optimizations. Their modest inhibitory activities versus KasA at a single compound concentration of 25 µM in contrast to their MIC values ≤12 µM suggest that KasA may not be their primary target within *M. tuberculosis*. This result highlights a significant challenge to the field regarding the prediction of the primary target of a small molecule. A previous effort in our labs with *M. tuberculosis* InhA also encountered this challenge^67^. We hypothesize that compound selection may be further augmented by machine learning models leveraging MIC shifts for compounds against a target overexpressor or knockdown strain as compared to the wild type strain. Significant precedent exists for the increase in a compound’s MIC when its protein target is overexpressed within the bacterium and likewise decreased when its target is knocked down within the bacterium^68^. Furthermore, we reiterate that these twelve hits are potent whole-cell active compounds with promising oral exposure in mice. Thus, they represent valuable starting points for new KasA inhibitors if their X-ray crystal structure with KasA can be determined to initiate a structure-based optimization^18^. Conversely, if a primary target is demonstrated to not be KasA and if an X-ray crystal structure of the primary target protein-inhibitor complex is attained, the same structure-based approach could also be successful. In either case, the promising values of MIC and PK AUC_0-5h_ of the hit compounds provide favorable starting points for optimization with the goal of delivering *in vivo* efficacy in an acute model of *M. tuberculosis* infection where JSF-5071 did not show promise.

These efforts to combine target engagement and molecule property prediction join other efforts in the field. In our laboratories alone, we have previously published on filtering high-throughput docking hits to *M. tuberculosis* InhA with an ML model for MIC^67^, predicting antitubercular whole-cell actives by predicting both MIC and Vero cell cytotoxicity CC_50_^69^, and predicting *Plasmodium berghei* growth inhibition and HepG2 cell cytotoxicity^70^.

In conclusion, we have presented an approach to combining high-throughput docking and molecular property ML models to afford 12 small molecule antituberculars with promising MIC and PK metrics. While the antitubercular hits offer varying levels of inhibition of the docking target KasA, they represent opportunities for hit-to-lead programs to optimize their target-based inhibition while further enhancing their whole-cell activity and PK profiles. We anticipate future opportunities to expand this approach to the multi-parameter optimization problem that is inherent to drug discovery and development.

## MATERIALS AND METHODS

### Computation

#### Docking and Structural Filtering

Docking was performed with QVina2^27^ via pyscreener^71^ where the X-ray crystal structure of *M. tuberculosis* KasA with JSF-3285 bound was used (PDB: 6P9L) as the target^56^ and compounds from the 2.1M Enamine HTS Collection (2020 release; www.enamine.net) were utilized as the ligands. The docking of the compound library was performed in batches using the Python package MolPAL^72^ with a batch size of 0.4% of the total library (ca. 8.2k compounds) using one random initialization batch and then five acquisition batches for a total of ∼51k docking evaluations. The bounding box was set to be [13:4; 16:2; 14:5] while retrieving a highly scored docked pose aligned with the experimental pose of JSF-3285 (root-mean-square distance of 0.765 Å) by setting the center to be [49:8; −7:9; −18:5]. The upper confidence bound acquisition score was used as the acquisition function. The optimization was run for five acquisition iterations. Input and configuration files may be found in our GitHub repository https://github.com/coleygroup/ml-for-tb/tree/main/data/2_molpal. The top 1,000 compounds based on the score of their best sampled pose were selected for further processing (Supplementary File S1).

A filtering process was performed to avoid candidates with undesired properties. Compounds with more than four rings were removed from the top 1,000 candidates to avoid the potential for DNA intercalation^73^. The nitro group, while present in various therapeutics, represents a mutagenicity/genotoxicity concern^74, 75^. Therefore, compounds with nitro groups were filtered out via SMARTS pattern, [$([NX3](=O)=O), $([NX3+](=O)[O- ])][!#8]. After these filters, 956 compounds remained (Supplementary File S1).

#### Machine Learning Property-Based Models

##### PK Dataset

The dataset used to model compound PK AUC_0-5h_, obtained using the Novartis snapshot PK protocol^76, 77^, consisted of 190 unique compounds. Each compound was orally administered via a 25 mg/kg oral dose with a formulation of 5% DMA/60% polyethylene glycol 300/35% (5% dextrose in water) to two female CD-1 mice. Blood samples were drawn 4 times over a 5 h period to enable the quantification of compound level within the mouse plasma. These data were subsequently used to calculate the area under the average plasma concentration versus time curve (AUC_0–5h_). The recorded AUC_0-5h_ ranged from 0 to 110,845 h*ng/mL, and the distribution was heavily skewed towards the minimum value. The samples were nearly equally divided (50.5% positives) around the 1,000 h*ng/mL mark, which in our experience has shown to be a reliable threshold for this metric for a hit compound^78^. Given these conditions, we tested an initial set of regression models, which revealed large confidence intervals for predictions as we expected. We, hence, reformulated the problem as a binary classification task using compounds with AUC_0-5h_ ≥ 1,000 h*ng/mL as positives and negatives as AUC_0-5h_ < 1,000 h*ng/mL.

##### MIC Datasets

The three datasets used for modeling compound MIC versus *M. tuberculosis* are publicly available datasets from high-throughput, whole-cell phenotypic screens of large chemical libraries by the Tuberculosis Antimicrobial Acquisition and Coordinating Facility (TAACF). The first effort involved 100,997 compounds, purchased from the ChemBridge Corporation and selected for diversity and drug-likeness, screened against the *M. tuberculosis* H37Rv strain in a single-dose format at 10 μg/mL^79^. Compounds with more than 90% inhibition were categorized as hits. The second effort involved 215,110 compounds from the NIH Small Molecule Repository developed through the Molecular Libraries Screening Center Network initiative^80^. There were minor changes to the screening protocol as compared to the Chembridge screen, including the dose concentration being set to 10 μM. The third effort focused on 25,671 kinase-specific inhibitors supplied by Life Chemicals, Inc., screened at a concentration of 10 μg/mL^81^. We refer to these three datasets as TAACF-ChemBridge, MLSMR, and TAACF-Kinase, respectively. Considering that TAACF-ChemBridge and TAACF-Kinase share the same screening concentration of 10 μg/mL, we combined and de-duplicated these to form a larger dataset which we refer to as the TAACF dataset. After merging and curation, the sizes of the TAACF and MLSMR datasets were 124,094 and 214,506, respectively.

##### Data Curation

Molecules with incomplete data and duplicates were removed. The RDKit package^82^ was used to generate the molecular SMILES strings, and duplicate compounds were removed by comparing the SMILES strings. Salts were stripped with the SaltRemover module in RDKit using a predefined list of SMARTS patterns. All datasets were split into training, validation, and test sets in a stratified manner, maintaining the same proportion of positive samples in each split. While the PK dataset contained a fairly equal number of positives and negatives (50.5% positives), the TAACF and MLSMR datasets had a high amount of class imbalance with 2.42% and 1.81% positives, respectively. We followed a commonly used train:validation:test split of 60:20:20 for the PK and TAACF datasets, but we used a 40:30:30 split for the MLSMR dataset. This is because in a highly imbalanced environment, a very small number of positives in the validation and test splits rendered the model evaluation unreliable.

##### Overview of Models

*Tree-based models* – The implementation of the Random Forest algorithm available in the scikit-learn package^85^ was utilized. XGBoost, LightGBM, and CatBoost were implemented using their own respective Python packages. *ChemBERTa-v2* – The ChemBERTa-v2 model is based on the RoBERTa architecture^88^ and has 5M, 10M, and 77M configurations, representing the size of the pretraining dataset. Each configuration was separately pretrained using two tasks: Masked Language Modeling and Multi-Task Regression (MTR). We used the 77M configuration consisting of three transformer blocks pretrained using MTR (’ChemBERTa-v2-MTR-77M’) as our feature extractor, which is reported to have competitive performance across datasets in the MoleculeNet benchmark^89^. To pretrain this configuration, the authors computed 200 deterministic molecular properties using RDKit and the model was trained to predict these properties simultaneously. We used the pretrained model checkpoint available in the Hugging Face Transformers API^90^. The extracted features were fed into a fully connected neural network, resulting in a binary output. The ChemBERTa-v2 feature extractor was finetuned during the training process. *Directed-MPNN* – We implemented the D-MPNN model using the Chemprop package^91^. The D-MPNN model’s feature extraction corresponds to its message passing phase (a series of message passing layers), and the classification head corresponds to its readout phase (a fully connected neural network). The best D-MPNN model for the MIC-MLSMR dataset used RDKit 2D features in addition to the molecular adjacency matrix. For other datasets, we did not see an improvement in performance after adding atom and bond features. All D-MPNN models were trained with an equal number of positives and negatives in each batch.

##### Feature Generation

Traditional models such as Random Forest and XGBoost used the 2D descriptors from the Molecular Operating Environment (MOE software version 2022.02) and Morgan fingerprints^83^ from the RDKit package as our starting set of molecular features. The D-MPNN model primarily used the adjacency matrix of the molecular graph, and in some instances also used the aforementioned fingerprints as atom features in conjunction with the adjacency matrix. Pretrained models such as ChemBERTa-v2 used specific tokenizers made publicly available by their creators that reduce a SMILES string to its component chemical tokens and assign integer token identifiers. We used the ChemBERTa-v2 tokenizer which has a vocabulary size of 591 and is based on Byte-Pair Encoding^84^. Token embeddings were learned during model training. The embedding dimension was determined through hyperparameter optimization.

##### Feature Engineering

We developed several pipelines (described in the Additional Computational Details of the Supplementary Materials) to reduce the feature space when creating the models. Our results showed that a “one size fits all” approach did not work best across datasets, and so we restricted each dataset to two promising pipelines each. All pipelines, however, used some if not all of the components outlined below. *Multicollinearity Handling* – We calculated pairwise feature correlation by determining both the Pearson’s correlation coefficient (ρ) and the R^2^ coefficient matrix. In general, we used Corr to denote the upper triangular region of the respective coefficient matrix, where Corr_i,j_ was the correlation between features X_i_ and X_j_. We removed the set of features {i | ∋j : Corr_i,j_ > t_corr_} from the full set of features where t_corr_ is a threshold specific to its pipeline. *Variance Thresholding* – This involved removing all features X_i_ having variance below a given threshold {t_var_: i | Var[X_i_] < t_var_}. *Importance Score* – To ensure that the features were useful in training, we filtered by feature importance. To achieve this, we trained k = 5 XGBoost models (denoted below by M_i_ where I ∊ {1,…,K}) in a k-fold fashion, leaving out a different 1/K of the training set each time and calculated an importance score. Features with zero importance score, {I | Importance-Score(X_i_) = 0} were removed. The importance score was calculated as follows, where X_i_ denotes the i-th feature vector (eqn. 1).

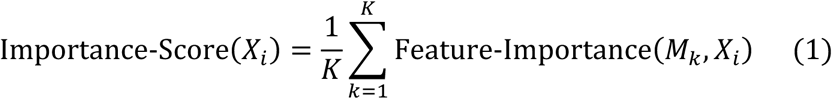

*Normalization* – Each of the MOE 2D descriptors was normalized by subtracting the mean and scaling them to unit variance. The means and variances of the features were stored and re-used at inference time to scale the features of the inference set. Normalization of the feature vector X_i_ was achieved as follows (eqn. 2):

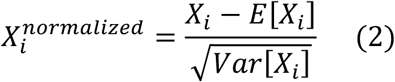

Further details regarding model training, model evaluation, and compound nomination may be found in the Additional Computational Details of the Supplementary Materials. Also, the datasets and code for the machine learning models presented herein may be found at our GitHub repository at https://github.com/coleygroup/ml-for-tb/tree/main/data/0_raw and https://github.com/coleygroup/ml-for-tb/tree/main/models.

### Chemistry

#### General Methods

All reagents were purchased from commercial suppliers and used without further purification unless noted otherwise. All chemical reactions occurring solely in an anhydrous organic solvent were carried out under an inert atmosphere of nitrogen or argon unless noted otherwise. Reactions performed at rt were typically at 21 – 24 °C. Analytical TLC was performed with Merck silica gel 60 F_254_ plates. Silica gel column chromatography was conducted with Teledyne Isco CombiFlash Companion or Rf+ systems. ^1^H NMR spectra were acquired on Bruker 500 and 600 MHz instruments and are listed in parts per million downfield from TMS. ^1^H NMR and ^13^C NMR spectra for JSF-5077 and JSF-5088 were acquired on a Bruker 500 MHz instrument at 105 ℃. Liquid chromatography-mass spectrometry (LC-MS) was performed on an Agilent 1260 HPLC coupled to an Agilent 6120 MS. All synthesized compounds were at least 95% pure as judged by their HPLC trace at 250 nm and were characterized by the expected parent ion/s in the MS. HRMS was performed on an Agilent 6230B Accurate Mass TOF MS.

#### Synthesis of 6-bromo-2-phenyl-N-(4-(pyridin-2-yl)thiazol-2-yl)quinoline-4-carboxamide (JSF-5071)

Synthesis of 6-bromo-2-phenylquinoline-4-carboxylic acid: To a solution of 5-bromoisatin (1.00 g, 4.43 mmol) in EtOH (15 mL) was added 33% KOH_(aq)_ (10 mL) and acetophenone (797 mg, 6.65 mmol, 1.5 equiv) and the reaction mixture was stirred at reflux for 72 h. The progress of the reaction was monitored by LC-MS. The solvent was removed under reduced pressure and the remaining aqueous layer was neutralized with 1N HCl_(aq)_ to pH 3. The precipitated solid was filtered, washed with water, and dried under the air to afford the desired compound as a yellow solid (700 mg, 2.14 mmol, 47.1%): ^1^H NMR (500 MHz, d_6_-DMSO) δ 14.1 (s, 1H), 8.93 (s, 1H), 8.51 (s, 1H), 8.26 (d, *J* = 7.2 Hz, 2H), 8.06 (d, *J* = 8.8 Hz, 1H), 7.94 (d, *J* = 8.7 Hz, 1H), 7.55 (m, 3H). Calculated for C_16_H_11_NBrO_2_ [M+H]^+^: 328.0, found 328.0.

To a stirred solution of 6-bromo-2-phenylquinoline-4-carboxylic acid (500 mg, 1.52 mmol) in DMF (10 mL) was added HATU (866 mg, 2.28 mmol, 1.5 equiv). The reaction mixture was stirred for 15 min. Then, DIPEA (0.80 mL, 4.6 mmol, 3.0 equiv) and 4-(pyridin-2-yl)thiazol-2-amine (269 mg, 1.52 mmol) were added. The reaction mixture was stirred at rt for 4 h. The progress of the reaction was monitored by TLC and LC-MS. Upon completion, the reaction mixture was subjected to the addition of ice-cold water and extracted with ethyl acetate (2 x 50 mL). The combined ethyl acetate extracts were washed with cold water and saturated aqueous brine solution, dried over anhydrous Na_2_SO_4_ and evaporated under reduced pressure to obtain a light brown crude liquid. The reaction product was purified via flash chromatography over silica gel eluting with MeOH (with 2% v/v 28 – 30% NH_4_OH)/DCM (0 – 10%) to obtain the title compound as an off-white solid (380 mg, 0.782 mmol, 51.1%): ^1^H NMR (500 MHz, d_6_-DMSO) δ 13.4 (s, 1H), 8.64 (d, *J* = 4.7 Hz, 1H), 8.62 (s, 1H), 8.55 (s, 1H), 8.40 (d, *J* = 7.6 Hz, 2H), 8.13 (d, *J* = 8.9 Hz, 1H), 8.02 (m, 3H), 7.91 (t, *J* = 7.8 Hz, 1H), 7.60 (m, 3H), 7.37 (t, *J* = 6.0 Hz, 1H). Also noted 3.3 (s, H_2_O), 1.27 (m), 0.8 (m). ^13^C NMR (126 MHz, d_6_-DMSO) δ 164.8, 158.1, 156.4, 151.9, 149.6, 149.4, 146.7, 138.0, 137.7, 137.4, 133.4, 131.8, 130.3, 128.9, 127.5, 127.2, 124.3, 122.9, 120.9, 120.1, 119.5, 112.6. Calculated for C_24_H_16_BrN_4_OS [M+H]^+^ = 487.0228, found 487.0238.

#### General Procedure A: Synthesis of N-(benzo[d][1,3]dioxol-5-yl)-1-(2-(4-fluorophenyl)quinoline-4-carbonyl)piperidine-4-carboxamide (JSF-5077)

Synthesis of *tert*-butyl 4-(benzo[d][1,3]dioxol-5-ylcarbamoyl)piperidine-1-carboxylate: To a stirred solution of 1-(*tert*-butoxycarbonyl)piperidine-4-carboxylic acid (300 mg, 2.18 mmol) in DMF (10 mL) was added HATU (1.25 g, 3.27 mmol, 1.5 equiv). The reaction mixture was stirred for 15 min. Then, DIPEA (1.14 mL, 6.54 mmol, 3.0 equiv) and benzo[d][1,3]dioxol-5-amine (551 mg, 2.40 mmol, 1.1 equiv) were added. The reaction mixture was stirred at rt for 4 h. The progress of the reaction was monitored by LC-MS. Upon completion, the reaction mixture subjected to the addition of ice-cold water and extracted with ethyl acetate (2 x 50 mL). The combined ethyl acetate extracts were washed with cold water and saturated aqueous brine solution, dried over anhydrous Na_2_SO_4_ and evaporated under reduced pressure to obtain a black liquid. The reaction product was purified via flash chromatography over silica gel eluting with ethyl acetate/hexane (0 to 60%) to obtain the desired product as a light brown liquid (450 mg, 1.29 mmol, 98.5%): ^1^H NMR (500 MHz, d_6_-DMSO) δ 9.80 (s, 1H), 7.30 (d, J = 2.1 Hz, 1H), 6.96 (dd, J = 8.4, 2.1 Hz, 1H), 6.82 (d, J = 8.4 Hz, 1H), 5.96 (s, 2H), 3.98 (m, 2H), 2.75 (s, 2H), 2.44 (m, 1H), 1.74 (m, 2H), 1.47 (m, 2H), 1.40 (s, 9H). Calculated for C_14_H_17_N_2_O_5_ [M+H-56]^+^: 293.1, found 293.1.

Synthesis of N-(benzo[d][1,3]dioxol-5-yl)piperidine-4-carboxamide: To a stirred solution of *tert*-butyl 4-(benzo[d][1,3]dioxol-5-ylcarbamoyl)piperidine-1-carboxylate (450 mg, 1.29 mmol, 1.0 equiv) in DCM (10 mL) at rt was added TFA (2 mL) and reaction mixture was stirred for 6 h. The progress of the reaction was monitored LC-MS. Upon completion, the reaction solvent was evaporated under reduced pressure to afford a light black residue. The residue was neutralized with a saturated aqueous solution of NaHCO_3_ and extracted with DCM (2 x 40 mL). The organic extracts were washed with saturated aqueous brine solution, dried over anhydrous Na_2_SO_4_ and evaporated under reduced pressure to obtain a light brown crude liquid (198 mg, 0.798 mmol, 61.8%): ^1^H NMR (500 MHz, d_6_-DMSO) δ 9.96 (s, 1H), 8.33 (s, 1H), 7.30 (s, 1H), 6.97 (d, J = 8.2 Hz, 1H), 6.83 (d, J = 8.4 Hz, 1H), 5.96 (s, 2H), 3.30 (d, J = 13.2 Hz, 2H), 2.87 (t, J = 11.4 Hz, 2H), 2.58 (m, 1H), 1.88 (m, 2H), 1.75 (m, 2H). Calculated for C_13_H_17_N_2_O_3_ [M+H]^+^: 249.1, found 249.1. The product was used as such without any further purification.

Synthesis of methyl 2-(4-fluorophenyl)quinoline-4-carboxylate: To a mixture of methyl 2-chloroquinoline-4-carboxylate (200 mg, 0.905 mmol) in dioxane/H_2_O (4:1 mL) was added Pd(PPh_3_)_4_ (52 mg, 0.045 mmol, 0.05 equiv). The reaction mixture was sparged with nitrogen for 15 min followed by the addition of (4-fluorophenyl)boronic acid (152 mg, 1.09 mmol, 1.2 equiv). Na_2_CO_3_ (288 mg, 2.72 mmol, 3.0 equiv) was added and again the reaction mixture was sparged with nitrogen for 15 min and stirred at 95 ℃ for 24 h. The progress of the reaction was monitored by LC-MS. The reaction mixture was cooled, subjected to the addition of 30 mL H_2_O, and extracted with ethyl acetate (2 x 30 mL). The combined organic layers were washed with saturated aqueous brine solution, dried over anhydrous Na_2_SO_4_ and evaporated under reduced pressure to obtain a light black liquid. The reaction product was purified via flash chromatography on silica gel eluting with ethyl acetate/hexane (0 to 60%) to obtain the desired product as an off-white solid (238 mg, 0.846 mmol, 93.7%): ^1^H NMR (500 MHz, d_6_-DMSO) δ 8.55 (dd, *J* = 8.5, 1.4 Hz, 1H), 8.48 (s, 1H), 8.37 (m, 2H), 8.17 (m, 1H), 7.87 (ddd, *J* = 8.4, 6.8, 1.4 Hz, 1H), 7.72 (ddd, *J* = 8.4, 6.8, 1.3 Hz, 1H), 7.42 (m, 2H), 4.03 (s, 3H). Also observed 3.32 (s, H_2_O). Calculated for C_17_H_13_FNO_2_ [M+H]^+^: 282.1, found 282.1.

Synthesis of 2-(4-fluorophenyl)quinoline-4-carboxylic acid: To a solution of methyl 2-(4-fluorophenyl)quinoline-4-carboxylate (238 mg, 0.846 mmol) in THF/MeOH (4:1, 10 mL) was added 2N NaOH_(aq)_ (4 mL) and the mixture was stirred at 70 ℃ for 5 h. The progress of the reaction was monitored by LC-MS. After completion, the solvent was evaporated under reduced pressure and the aqueous layer was adjusted with aqueous 1N HCl_(aq)_ to pH 3 – 4. The precipitated solid was filtered and dried under the air to afford the desired compound as a light yellow solid (213 mg, 0.797 mmol, 94.1%): ^1^H NMR (500 MHz, d_6_-DMSO) δ 14.0 (s, 1H), 8.64 (d, *J* = 8.5 Hz, 1H), 8.40 (d, *J* = 47.4 Hz, 3H), 8.15 (d, *J* = 8.5 Hz, 1H), 7.85 (t, *J* = 7.9 Hz, 1H), 7.71 (d, *J* = 7.9 Hz, 1H), 7.39 (t, *J* = 8.8 Hz, 2H). Calculated for C_16_H_11_FNO_2_ [M+H]^+^: 268.1, found 268.1.

To a stirred solution of 2-(4-fluorophenyl)quinoline-4-carboxylic acid (213 mg, 0.797 mmol) in DMF (10 mL) was added HATU (454 mg, 1.20 mmol, 1.5 equiv). The reaction mixture was stirred for 15 min. Then, DIPEA (0.41 mL, 2.39 mmol, 3.0 equiv) and N- (benzo[d][1,3]dioxol-5-yl)piperidine-4-carboxamide (198 mg, 0.797 mmol, 1.0 equiv) were added. The reaction mixture was stirred at rt for 4 h. The progress of the reaction was monitored by LC-MS. Upon completion, the reaction mixture subjected to the addition of ice-cold water and extracted with ethyl acetate (2 x 50 mL). The combined ethyl acetate extracts were washed with cold water and saturated aqueous brine solution, dried over anhydrous Na_2_SO_4_ and evaporated under reduced pressure to obtain a light black liquid. The reaction product was purified via flash chromatography over silica gel eluting with ethyl acetate/hexane (0 to 90%) to obtain the title product as a light brown solid (100 mg, 0.201 mmol, 25.2%): ^1^H NMR (500 MHz, d_6_-DMSO, 105 °C) δ 9.45 (s, 1H), 8.35 (m, 2H), 8.13 (d, J = 8.9 Hz, 1H), 8.02 (s, 1H), 7.83 (m, 2H), 7.66 (t, J = 7.5 Hz, 1H), 7.35 (t, J = 8.9 Hz, 2H), 7.25 (d, J = 2.1 Hz, 1H), 6.97 (dd, J = 8.4, 2.1 Hz, 1H), 6.79 (d, J = 8.3 Hz, 1H), 5.94 (s, 2H), 4.67 (s, 1H), 3.41 (s, 1H), 3.13 (t, J = 12.4 Hz, 2H), 2.65 (m, 1H), 2.02 (m, 1H), 1.65 (m, 3H). ^13^C NMR (126 MHz, d_6_-DMSO, 105 °C) δ 171.7, 165.3, 161.9 (*J_CF_* = 255.8 Hz), 154.5, 147.3, 146.6, 143.5, 142.5, 134.4 (*J_CF_* = 2.5 Hz), 133.1, 129.8, 129.1, 129.0 (*J_CF_* = 16.7 Hz), 126.7, 124.0, 122.4, 114.9 (*J_CF_* = 21.4 Hz), 114.6, 112.1, 107.2, 101.5, 100.3, 41.7. Two carbons (i.e., most likely piperidine carbons) were not observed. Calculated for C_29_H_25_FN_3_O_4_ [M+H]^+^: 498.1829, found 498.1807.

#### Synthesis of N-(benzo[d][1,3]dioxol-5-yl)-1-(2-(p-tolyl)quinoline-4-carbonyl)piperidine-4-carboxamide (JSF-5088) was followed as per General Procedure A for JSF-5077)

^1^H NMR (500 MHz, d_6_-DMSO, 105 °C): δ 9.52 (s, 1H), 8.20 (d, J = 8.2 Hz, 2H), 8.13 (m, 1H), 8.00 (s, 1H), 7.80 (m, 2H), 7.64 (m, 1H), 7.37 (d, J = 7.9 Hz, 2H), 7.26 (d, J = 2.0 Hz, 1H), 6.97 (dd, J = 8.4, 2.1 Hz, 1H), 6.79 (d, J = 8.4 Hz, 1H), 5.94 (s, 2H), 4.68 (s, 1H), 3.39 (s, 1H), 3.11 (t, J = 12.5 Hz, 2H), 2.64 (m, 1H), 2.42 (s, 3H), 2.02 (m, 1H), 1.70 (m, 3H). Also observed 3.03 (s, H_2_O); ^13^C NMR (126 MHz, d_6_-DMSO, 105 °C) δ 171.7, 165.4, 155.6, 147.4, 146.6, 143.2, 142.5, 138.9, 135.2, 133.1, 129.6, 129.1, 128.8, 126.7, 126.4, 124.0, 122.4, 114.5, 112.1, 107.2, 101.5, 100.3, 41.7, 20.2. Two carbons (i.e., most likely piperidine carbons) were not observed. Calculated for C_30_H_28_N_3_O_4_ [M+H]^+^: 494.2080, found 494.2085.

#### Synthesis of 2-oxo-N-(4-(N-(3-(trifluoromethyl)phenyl)sulfamoyl)phenyl)-2H-chromene-3-carboxamide (JSF-5089)

Synthesis of 4-nitro-N-(3-(trifluoromethyl)phenyl)benzenesulfonamide: To a stirred solution of 3- (trifluoromethyl)aniline (1.0 g, 6.2 mmol) in DCM (20 mL) was added pyridine (1.00 mL, 12.4 mmol, 2.0 equiv) and 4-nitrobenzenesulfonyl chloride (1.51 g, 6.83 mmol, 1.1 equiv). The reaction mixture was stirred at rt for 15 h. The progress of the reaction was monitored by LC-MS. Upon completion, a saturated aqueous solution of NaHCO_3_ was added to the reaction mixture which was then extracted with DCM (2 x 80 mL). The combined DCM extracts were washed with saturated aqueous brine solution, dried over anhydrous Na_2_SO_4_ and evaporated under reduced pressure to obtain an off-white solid. The reaction product was purified via flash chromatography over silica gel eluting with ethyl acetate/hexane (0 to 60%) to obtain the desired product as a white solid (2.0 g, 5.7 mmol, 93%): ^1^H NMR (500 MHz, d_6_-DMSO) δ 11.1 (s, 1H), 8.39 (d, *J* = 8.9 Hz, 2H), 8.03 (d, *J* = 8.9 Hz, 2H), 7.52 (t, *J* = 8.3 Hz, 1H), 7.41 (m, 3H). Calculated for C_13_H_10_F_3_N_2_O_4_S [M+H]^+^: 347.0, found 347.0.

Synthesis of 4-amino-N-(3-(trifluoromethyl)phenyl)benzenesulfonamide: To a solution of 4-nitro- N-(3-(trifluoromethyl)phenyl)benzenesulfonamide (500 mg, 1.45 mmol) in MeOH (20 mL) at rt was added Zn (1.41 g, 21.7 mmol, 15 equiv) and NH_4_Cl (1.15 g, 21.7 mmol, 15 equiv). The reaction mixture was stirred at rt for 3 h. The progress of reaction was monitored via LC-MS. The reaction mixture was filtered through a bed of Celite which was washed with MeOH (2 x 80 mL). The volatiles were evaporated under reduced pressure to obtain an off-white solid. To the solid was added water (30 mL) and the mixture was extracted with 5% MeOH/EtOAc (2 x 60 mL). The organic layer was washed with a saturated aqueous brine solution, dried over anhydrous sodium sulfate and then concentrated *in vacuo* to obtain the desired product as a sticky colorless liquid (450 mg, 1.42 mmol, 98.2%): ^1^H NMR (500 MHz, d_6_-DMSO) δ 10.3 (s, 1H), 7.45 (t, *J* = 7.9 Hz, 1H), 7.41 (d, *J* = 8.8 Hz, 2H), 7.33 (m, 3H), 6.54 (d, *J* = 8.8 Hz, 2H), 6.03 (s, 2H). Also noted 4.04 (q, EtOAc), 2.00 (s, EtOAc), and 1.17 (t, EtOAc). Calculated for C_13_H_12_F_3_N_2_O_4_S [M+H]^+^: 317.1, found 317.0. The product was used as such for the next step without any further purification.

To a stirred solution of 2-oxo-2H-chromene-3-carboxylic acid (282 mg, 1.42 mmol) in DMF (10 mL) was added HATU (811 mg, 2.13 mmol, 1.5 equiv). The reaction mixture was stirred for 15 min. Then, DIPEA (0.73 mL, 4.2 mmol, 3.0 equiv) and 4-amino-N-(3- (trifluoromethyl)phenyl)benzenesulfonamide (450 mg, 1.42 mmol, 1.0 equiv) were added. The reaction mixture was stirred at rt for 4 h. The progress of the reaction was monitored by LC-MS. Upon completion, the reaction mixture was subjected to the addition of ice-cold water and extracted with ethyl acetate (3 x 60 mL). The combined ethyl acetate extracts were washed with cold water and saturated aqueous brine solution, dried over anhydrous Na_2_SO_4_ and evaporated under reduced pressure to obtain a light brown liquid. The reaction product was purified via flash chromatography over silica gel eluting with MeOH/DCM (0 to 10%) to obtain a yellow solid. The solid was suspended in ethanol and filtered to afford the title compound as a light yellow solid (80 mg, 0.16 mmol, 11%): ^1^H NMR (500 MHz, d_6_-DMSO): δ 10.9 (s, 1H), 10.7 (s, 1H), 8.89 (s, 1H), 8.00 (dd, J = 7.9, 1.7 Hz, 1H), 7.90 (d, J = 8.5 Hz, 2H), 7.80 (m, 3H), 7.55 (d, J = 8.4 Hz, 1H), 7.48 (dt, J = 11.8, 7.7 Hz, 2H), 7.38 (m, 3H). Also noted 3.3 (s, H_2_O). ^13^C NMR (151 MHz, d_6_-DMSO) δ 160.6, 160.1, 153.9, 147.70, 142.1, 138.7, 134.5, 133.7, 130.6, 130.8, 129.5 (*J_CF_* = 31.7 Hz), 128.2, 125.3, 123.2, 122.9 (*J_CF_* = 271.8 Hz), 120.3 (*J_CF_* = 4.5 Hz), 119.9, 119.9, 118.4, 116.3, 115.5 (*J_CF_* = 4.5 Hz). Calculated for C_23_H_16_F_3_N_2_O_5_S [M+H]^+^: 489.0732, found 489.0729.

### Biological Assays

#### KasA Binding Assay

Prior to labeling, KasA (obtained via our published protocol^92^) was diluted from 120 μM to 20 μM in Buffer A (500 mM NaCl, 20 mM CHES pH 9.5). The RED-NHS labeling kit (NanoTemper Technologies) was then used to label KasA. Labeled KasA was exchanged into 10 mM HEPES pH 7.4 and 150 mM NaCl (Buffer B) and unincorporated label removed using a PD-10 column. 100 nM working stocks of labeled KasA were made in Buffer B supplemented with 0.2% Pluronic F-127. Stocks of the inhibitor were made by dissolving the compound in DMSO. Inhibitor was then serially diluted into individual Eppendorf tubes (final [DMSO] = 5% in each tube) and incubated with equal volumes of 100 nM labeled KasA (final [KasA] = 50 nM). This mixture was incubated in the dark for 30 min at rt. After incubation, the samples were transferred into Premium Coated Capillaries (NanoTemper Technologies) and analyzed using a Monolith NT.115 Nano-RED Instrument set to 40% LED and 40% MST power. Binding affinity was calculated from duplicate data analyzed by the MO.Affinity Analysis software v2.3 (NanoTemper Technologies). When significant binding was detected in the binding affinity measurements, i.e., signal:noise greater than 8, the experiments were carried out in duplicate.

#### KasA Functional Inhibition Assay

Protein overexpression and purification: *M. tuberculosis* KasA was overexpressed and purified as previously described^92^. A 6xHis-SUMO (small ubiquitin-related modifier sequence optimized for *E. coli* overexpression) fusion *M. tuberculosis* MabA gene was purchased from Synbio Technologies. MabA was cloned into a pET28a plasmid between restriction sites NdeI and XhoI which was then transformed into *E. coli* T7 Express *lysY/I^q^* competent cells (New England Biolabs). His-SUMO-MabA was overexpressed in *E. coli* grown in LB medium containing 50 μg/mL kanamycin until OD_600_ = 0.70 ∼ 0.75 at 37 °C. Expression was induced with 300 μM IPTG and cells were grown for 3 h at 37 °C, followed by centrifugation at 5,000 rpm for 15 min. MabA was purified using a standard protocol for His-SUMO fusion constructs^93^. The cell pellet was resuspended in Buffer A [50 mM Tris, pH 8, 150 mM NaCl, 10% glycerol (v/v)] along with 1 mM phenylmethylsufonyl fluoride (PMSF), cocktail cOmplete Mini EDTA-free protease inhibitor cocktail (Roche), and 1 mM tris(2-carboxyethyl)phosphine (TCEP) and lysed by sonication. Lysate was separated from cell debris by centrifugation at 18,000 rpm for 30 min at 4 °C. An initial Ni-NTA affinity purification step was performed to isolate His-SUMO-MabA, where Ni-NTA resin equilibrated with buffer A containing 20 mM imidazole was incubated with the clarified lysate for 30 min at 4 °C with rotation, followed by elution with increasing concentrations of imidazole. Next, the purified His-SUMO-MabA was cleaved by His-SUMO protease (purchased from MCLAB) to give native MabA and His-SUMO. For the cleavage reaction, the pooled fractions were diluted to 1 mg/mL in buffer A plus 0.2% NP-40 and incubated with 1 unit SUMO protease per 2 μg His-SUMO-MabA substrate for 1 h at 30 °C. The reaction product was buffer exchanged via PD-10 desalting column into 50 mM Tris, pH 8. Finally, MabA was isolated via another Ni-NTA chromatography step: the buffer exchanged product was incubated with fresh Ni-NTA resin for 30 min at 4 °C with 5 mM imidazole. The unbound fraction containing MabA was collected and dialyzed into buffer containing 50 mM Tris, pH 8, 10% glycerol, 25 mM NaCl, and 1 mM TCEP, flash frozen, and stored at −80 °C.

##### M. tuberculosis

AcpM was overexpressed and purified according to previously described protocols^47, 49, 94, 95^. AcpM was cloned into a pET28b plasmid via NdeI and XhoI restriction sites. The resulting N-terminal 6xHis-tagged AcpM pET28b vector was transformed and overexpressed in *E. coli* T7 Express *lysY/I^q^* competent cells. Cultures were grown at 37 °C in LB media with 50 μg/mL kanamycin until OD_600_ = 0.6 ∼ 0.7, and then expression was induced with 300 µM IPTG for 2.5 h at 37 °C. Cells were harvested by centrifugation at 5,000 rpm, 4 °C. Cell pellets were resuspended in buffer A containing 1 mM PMSF and cOmplete Mini EDTA-free protease inhibitor cocktail (Roche) and lysed by sonication. Insoluble protein and cell debris were separated by centrifugation at 18,000 rpm for 30 min at 4 °C. Clarified lysate was incubated with Ni-NTA resin equilibrated with buffer A for 1 h at 4 °C with rotation in the presence of 5 mM imidazole. His-AcpM was eluted with increasing concentrations of imidazole. Collected fractions were dialyzed overnight at 4 °C into buffer containing 50 mM Tris, pH 8, 20 mM NaCl, and 10% glycerol, followed by an additional 2 h dialysis. The dialysate was collected and centrifuged for 30 min at 18,000 rpm, 4 °C. Further purification of AcpM was performed via an AKTA Prime Plus FPLC with a Resource Q column (Global Life Sciences Solutions) in buffer 50 mM Tris, pH 8 and 10% glycerol, and eluted via a gradient of 0 to 400 mM NaCl over 100 mL (∼17 column volumes). Fractions containing acyl-AcpM were collected according to SDS-PAGE analysis, and LC-MS analysis revealed a mixture of of the acyl-AcpM and holo-AcpM species (Fig S5; LC-MS was performed by the Princeton University Proteomics & Mass Spectrometry Facility).

Enzyme Assays. Compound inhibition was determined via a functional assay coupling *M. tuberculosis* KasA activity to the next enzyme in the FAS-II pathway, MabA, similar to previously published approaches^46, 47, 49, 96^. Assays included Tween 20 at 1% of the critical micelle concentration to avoid non-specific inhibition of the enzyme due to aggregation^67, 97^. The coupled assay used 200 nM KasA, 5 μM MabA, 500 μM NADPH in a buffer containing 100 mM HEPES, pH 7, 1 mM TCEP, 0.5 μg/mL Tween 20, and 1.5% DMSO. Inhibitors were preincubated for 10 min at 25 °C before initiating the reaction with 50 μM of each substrate, the acyl-AcpM mixture (preparation described above) and malonyl-CoenzymeA(malonyl-CoA; MilliporeSigma). It has previously been shown that KasA can tolerate the CoA-bound malonyl substrate but prefers the acyl carrier protein (AcpM)-bound acyl substrate^96^. The assay followed the absorbance at 340 nm in a BioTek Synergy HTX 96-well plate reader over 10 min at 25 °C. Initial rates were determined from the raw data and adjusted based on background rates determined in the absence of KasA. Percent inhibition was calculated based on the initial rate with no inhibitor equal to 0%. The KasA functional assay demonstrated potent inhibition by JSF-3285 as a positive control (92.6 ± 3.5%). Furthermore, compounds were counter-screened against MabA. The MabA counter screen consisted of 500 nM MabA, 500 μM NADPH, and 500 μM acetoacetyl-CoA (AcAc-CoA; MilliporeSigma) in the same buffer as described above: 100 mM HEPES, pH 7, 1 mM TCEP, and 0.5 μg/mL Tween 20, 1.5% DMSO. Inhibitors were again preincubated for 10 min at 25 °C before initiating with AcAc-CoA and monitoring the reaction (λ*_max_*^NADPH^ = 340 nm) for 10 min at 25 °C.

#### Bacterial Strains and Media

##### M. tuberculosis

H37Rv and Erdman strains were from laboratory stocks and have been verified by whole-genome sequencing^11^. The strains were cultured in Middlebrook 7H9 media supplemented with 10% oleic acid-albumin-dextrose-catalase (OADC-Becton Dickinson, Sparks, MD), glycerol (0.2%) and 0.05% (w/v) Tween 80 in liquid media at 37 °C and shaking at 50 rpm. Middlebrook 7H11 agar (Becton Dickinson) supplemented with 0.5% glycerol (v/v) was used for growth on solid media at 37 °C.

#### Antibacterial Growth Inhibition Assays

The MIC assay was conducted in a 96-well microplate (Fisher #FB012932) with the microplate Alamar blue assay (MABA).^98^ Briefly, the test compounds were serially diluted in 100 µL of growth media (7H9-ADS – albumin-dextrose-sodium chloride) and 100 µL cultures (1:1000 dilution of OD_595_ = 0.2) were added to each well. The alamarBlue HS Cell Viability Reagent (Invitrogen #A50101) supplemented with 20% Tween 80 was added to each well after 7 d of incubation at 37 °C and the plates were further incubated for another 24 h to allow for the viability signal to develop (read at 570 nm absorbance and normalized to 600 nM as per manufacturer’s instructions).

#### Mammalian Cell Cytotoxicity Assay

Vero cells (African green monkey kidney epithelial cells; ATCC CCL-81) were cultured in a 96-well plate, at a concentration of 5,000 – 10,000 cells/well, and incubated for 2 – 3 h to allow the cells to settle. Test compounds were diluted separately in Dulbecco’s Modified Eagle Medium (DMEM) with 10% FBS to generate test concentrations typically ranging from 200 to 0.2 µM. The serial dilutions were then added to the plated cells and incubated for 48 – 72 h at 37 °C. The viability of Vero cells exposed to each compound was determined using the MTT [3-(4,5-dimethyl-2-thiazoyl)-2,5-diphenyl-2H-tetrazolium bromide] cell viability kit (Abcam ab211091). The CC_50_ was determined as the minimum test compound concentration to afford 50% growth inhibition of the model cell line.

#### Mouse Pharmacokinetics and Dose Tolerability Studies

Animal studies were carried out in accordance with the guide for the care and use of Laboratory Animals of the National Institutes of Health, with approval from the Institutional Animal Care and Use Committee (IACUC) of Hackensack Meridian Health. All animals were maintained under specific pathogen-free conditions and fed water and chow *ad libitum*, and all efforts were made to minimize suffering or discomfort. In the 5 h PK studies, two female CD-1 mice received a single dose of experimental compound administered orally at 25 mg/kg in 5% DMA/60% PEG300/35% D5W (5% dextrose in water), and blood samples were collected in K_2_EDTA coated tubes pre-dose, 0.5, 1, 3 and 5 h post-dose. In iv/po PK studies to determine oral bioavailability, groups of three female CD-1 mice received a single dose of experimental compound administered orally at 25 mg/kg in a 0.5% CMC/0.5% Tween 80 suspension or in a solution formulation of 1:9 DMSO:20% Solutol HS15, or intravenously at 5 mg/kg in 5% DMA/95% (4 % Cremophor EL). Blood samples were collected in K_2_EDTA coated tubes 0.25, 0.5, 1, 3, 5 and 8 h post-dose in the oral arm, and 0.033, 0.25, 0.5, 1 and 3 h post dose in the intravenous arm. Blood was kept on ice and centrifuged to recover plasma, which was stored at −80 °C until analysis by HPLC coupled to tandem mass spectrometry (LC-MS/MS). Oral bioavailability was reported as 100% multiplied by the dose-normalized plasma exposure of compound with oral dosing divided by the dose-normalized plasma exposure of compound with intravenous dosing. In the dose tolerability/proportionality study, three female CD-1 mice were dosed by oral gavage daily for 4 d with the compound in a solution formulation using 1:9 DMSO:20% Solutol HS15. Prior to dosing, the compound formulation was mixed and vortexed. The mice were weighed and observed daily. Their behavior, drinking and feeding patterns, and feces were monitored and recorded. Plasma samples were drawn on day 1 after compound administration at 0.5, 1, 3, 5, 8, and 24 h. Upon necropsy, liver, gallbladder, kidney and spleen pathology were observed for abnormalities.

LC-MS/MS analysis was performed on a Sciex Applied Biosystems Qtrap 6500+ triple-quadrupole mass spectrometer coupled to a Shimadzu Nexera X2 UHPLC system to quantify each drug in plasma, and chromatography was performed on an Agilent Zorbax SB-C8 column (2.1×30 mm; particle size, 3.5 µm) using a reverse phase gradient elution. Milli-Q deionized water with 0.1% formic acid (A) was utilized for the aqueous mobile phase and 0.1% formic acid in acetonitrile (B) for the organic mobile phase. The gradient was: 5-90% B over 2 min, 1 min at 90% B, followed by an immediate drop to 5% B and 1 min at 5% B. Multiple-reaction monitoring of parent/daughter transitions in electrospray positive-ionization mode was used to quantify all molecules. Sample analysis was accepted if the concentrations of the quality control samples and standards were within 20% of the nominal concentration. Data processing was performed using Analyst software (version 1.6.2; Applied Biosystems Sciex). Neat 1 mg/mL DMSO stocks for all compounds were first serial diluted in 50/50 acetonitrile/water and subsequently serial diluted in drug free CD-1 mouse plasma (K_2_EDTA, Bioreclamation IVT, NY) to create standard curves (linear regression with 1/x^2 weighting) and quality control (QC) spiking solutions. 20 µL of standards, QCs, control plasma, and study samples were extracted by adding 200 µL of acetonitrile/methanol 50/50 protein precipitation solvent containing the internal standard (10 ng/mL verapamil). Extracts were vortexed for 5 min and centrifuged at 4,000 rpm for 5 min. 100 µL of supernatant was transferred for HPLC-MS/MS analysis and diluted with 100 µL of Milli-Q deionized water. Plasma AUC_0-t_ was determined for each dosing group by trapezoidal integration.

#### Mouse Low-Dose Aerosol Acute Model of *M. tuberculosis* Infection Assay

On day 0, BALB/c mice (5 to 6-week-old females: weight range 18 to 21 g) were infected with an inoculum of *M. tuberculosis* Erdman mixed with 5 mL of phosphate-buffered saline (PBS) (4 x 10^6^ CFU/mL) using a Glas-Col whole-body aerosol unit.^58, 59^ An average lung infection of 1.94 log_10_ CFU per mouse was confirmed at 3 d post-infection (dpi) in n = 3 animals. Five mice per group were sacrificed at the start of treatment (7 dpi) and at 21 dpi after receiving JSF-5071 qd po (25 mg/kg; formulation of 1:9 DMSO:20% Solutol HS15), RIF bid po (10 mg/kg; formulation of 0.5% CMC/0.5% Tween 80), or the vehicle (0.5% CMC/0.5% Tween 80) qd po from 7 dpi through 18 dpi for a total of 12 consecutive days of dose administration. Due to limited resources and the previously mentioned formulation issues with JSF-5071, we were not able to match the vehicle control of 0.5% CMC/0.5% Tween 80 (which is the formulation used for the RIF treatment) with the formulation used for JSF-5071 (1:9 DMSO:20% Solutol HS15). Whole lungs were homogenized in 5 mL of PBS-Tween 80 (0.05%). CFU counts were determined by plating serial dilutions of homogenates onto Middlebrook 7H11 agar with OADC. Colonies were counted after at least 21 d of incubation at 37 °C. Ordinary one-way ANOVA with Tukey’s *post hoc* multiple comparisons test was used for statistical comparison with vehicle control and all individual treatment groups.

## AUTHOR INFORMATION

### Corresponding Authors

Joel S. Freundlich – Department of Pharmacology, Physiology, and Neuroscience and Division of Infectious Disease, Department of Medicine and the Ruy V. Lourenço Center for the Study of Emerging and Re-emerging Pathogens, Rutgers University - New Jersey Medical School, Newark, New Jersey USA;

Connor W. Coley – Department of Chemical Engineering, Department of Electrical Engineering and Computer Science, Massachusetts Institute of Technology, Cambridge, MA 02139

### Authors (Current Addresses are Noted)

Vedang Warapande – Department of Pharmacology, Physiology, and Neuroscience, Rutgers University – New Jersey Medical School, Newark, New Jersey, USA;

Fanwang Meng – Department of Chemistry, Queens University, Kingston, Ontario Canada, K7L 3N6;

Alexandra Bozan – Department of Pharmacology, Physiology, and Neuroscience, Rutgers University – New Jersey Medical School, Newark, New Jersey, USA;

David E. Graff – Atomwise, San Francisco, California;

Jenna C. Fromer – Department of Chemical Engineering, Massachusetts Institute of Technology, Cambridge, MA 02139, USA;

Khadija Mughal – Department of Pharmacology, Physiology, and Neuroscience, Rutgers University – New Jersey Medical School, Newark, New Jersey, USA;

Faheem K. Mohideen – Department of Pharmacology, Physiology, and Neuroscience, Rutgers University – New Jersey Medical School, Newark, New Jersey, USA;

Shivangi – Department of Pharmacology, Physiology, and Neuroscience, Rutgers University – New Jersey Medical School, Newark, New Jersey, USA;

Sindhuja Paruchuri – Center for Discovery & Innovation, Hackensack Meridian Health, Nutley, New Jersey, USA;

Melanie L. Johnston – Department of Chemistry and Biochemistry, Northern Arizona University, Flagstaff, Arizona, USA;

Pankaj Sharma – Department of Pharmacology, Physiology, and Neuroscience, Rutgers University – New Jersey Medical School, Newark, New Jersey, USA;

Timothy R. Crea – Department of Pharmacology, Physiology, and Neuroscience, Rutgers University – New Jersey Medical School, Newark, New Jersey, USA;

Reshma S. Rudraraju – Department of Microbiology, Biochemistry and Molecular Genetics, Rutgers University – New Jersey Medical School, Newark, New Jersey, USA;

Amir George – Department of Pharmacology, Physiology, and Neuroscience, Rutgers University – New Jersey Medical School, Newark, New Jersey, USA;

Camilla Folvar – Center for Discovery & Innovation, Hackensack Meridian Health, Nutley, New Jersey, USA;

Andrew M. Nelson – Center for Discovery & Innovation, Hackensack Meridian Health, Nutley, New Jersey, USA;

Matthew B. Neiditch – Department of Microbiology, Biochemistry and Molecular Genetics, Rutgers University – New Jersey Medical School, Newark, New Jersey, USA;

Matthew D. Zimmerman – Center for Discovery & Innovation, Hackensack Meridian Health, Nutley, New Jersey, USA;

## Supporting information

Molecular Formula Strings

## Author Contributions

The manuscript was written with contributions of all authors. All of the authors approved the final version of the manuscript.

## Notes

The authors declare no competing financial interests.

## ACKNOWLEDGEMENTS

This work was supported by NIH grants R21AI169342 to JSF and CWC and GM093854 to M.L.J..

## SUPPORTING INFORMATION

Performance of select models on the experimentally validated 93 compounds, evaluated retrospectively, Feature engineering and pipeline details for select models, KasA functional inhibition data for select compounds, MabA functional inhibition data for select compounds, Mouse PK parameters for JSF-5071 with a formulation of 5% DMA/95%(4% Cremophor) for iv dosing. (Tables S1 – S5) Counterscreen of twelve hits versus MabA, JSF-5071 mouse po and iv PK profile, Dose tolerability mouse PK study for JSF-5071 with data for day 4/4 dosing, Evaluation of JSF-5071 in a mouse acute model of *M. tuberculosis* infection with qd dosing, Characterization of isolated mixture of the acyl-AcpM and holo-AcpM species. (Figures S1 – S5). (PDF) Molecular formula strings (CSV). Supplementary File S1 (XLSX).

## ABBREVIATIONS USED

MIC: minimum inhibitory concentration
PK: pharmacokinetic
TB: tuberculosis
FAS-II: type Ⅱ fatty acid synthase
INH: isoniazid
area under the curve for 0 – 5 h: AUC_0-5h_
po: oral
MolPAL: Molecular Pool-based Active Learning
GNN: graph neural network
D-MPNN: directed message passing neural network
BCE: binary cross entropy
AUROC: area under the Receiver Operating Characteristic curve
AP: average precision
max. F1 score: maximum F1 score across all thresholds
TPE: Tree-Structured Parzen Estimator
CC_50_: the minimum compound concentration to afford 50% growth inhibition of this model mammalian cell line
iv: intravenous
MST: microscale thermophoresis
%F: oral bioavailability
qd: once-daily
bid: twice-daily
CFUs: colony-forming units
ADMET: Absorption-Distribution-Metabolism-Toxicity
MTR: Multi-Task Regression
LC-MS: liquid chromatography-mass spectrometry
LC-MS/MS: HPLC coupled to tandem mass spectrometry
dpi: days post-infection
PBS: phosphate-buffered saline.

## FOR TABLE OF CONTENTS ONLY

Source for graphic: NIAID Visual & Medical Arts. (10/7/2024). Funnel diagram. NIAID NIH BIOART Source. bioart.niaid.nih.gov/bioart/167

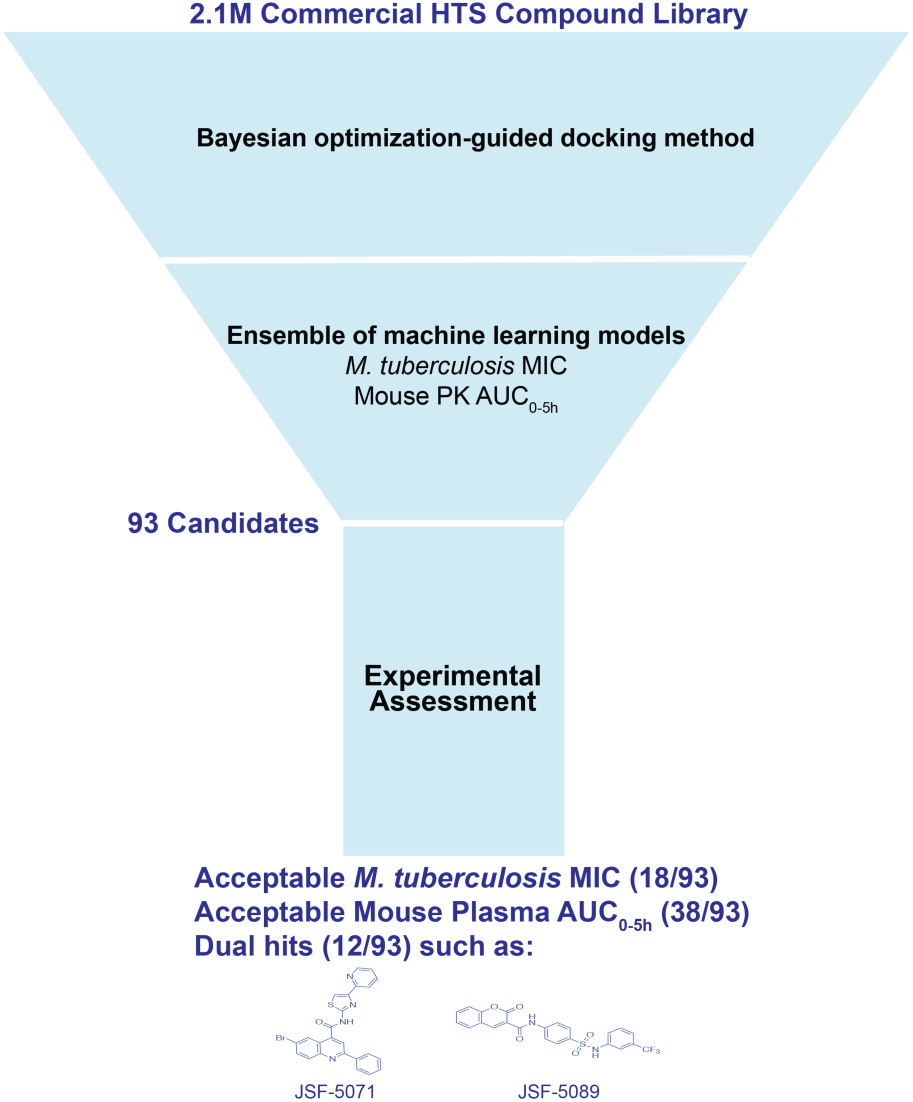

