## Supplemental Materials for "Identification of Antituberculars with Favorable Potency and Pharmacokinetics through Structure-Based and Ligand-Based Modeling"

**TABLE OF CONTENTS**


Figure S1. Counterscreen of twelve hits versus MabA. Page S3

Figure S2. JSF-5071 mouse po and iv PK profile. Page S4

Figure S3. Dose tolerability mouse PK study for JSF-5071 with data for day 4/4 dosing. Page S5

Figure S4. Evaluation of JSF-5071 in a mouse acute model of *M. tuberculosis* infection with qd dosing. Page S6

Figure S5. Characterization of isolated mixture of the acyl-AcpM and holo-AcpM species. Page S7

Table S1. Performance of select models on the experimentally validated 93 compounds, evaluated retrospectively. Page S8

Table S2. Feature engineering and pipeline details for select models. Page S9

Table S3. KasA functional inhibition data for select compounds. Page S10

Table S4. MabA functional inhibition data for select compounds. Page S11

Table S5. Mouse PK parameters for JSF-5071 with a formulation of 5% DMA/95%(4% Cremophor) for iv dosing. Page S12

Materials and Methods Page S13

Additional Computational Details Page S13

Additional Compound Characterization Data Page S15


Supplementary File S1.XLSX

Molecular formula strings (CSV) Please refer to Molecular_formula_strings.CSV

**Figure S1. Counterscreen of twelve hits versus MabA.** (A) MabA inhibition values for the 12 hit compounds (at 25 μM compound concentration) are depicted with the reported value representing the mean ± standard deviation from 6 replicates. (B) An attempted $IC_{50}$ curve (each data point representative of the mean ± standard deviation for two replicates) for Z1033301210 – the only compound to show >10% inhibition of MabA under the assay conditions in (A). Solubility constraints for Z1033301210 did not allow its assay above 100 μM. Conditions for (A) and (B): 25 °C, 500 nM MabA, 100 mM HEPES pH 7, 500 μM NADPH, 500 μM AcAcCoA, 1 mM TCEP, 1% CMC Tween-20, 1.5% DMSO.

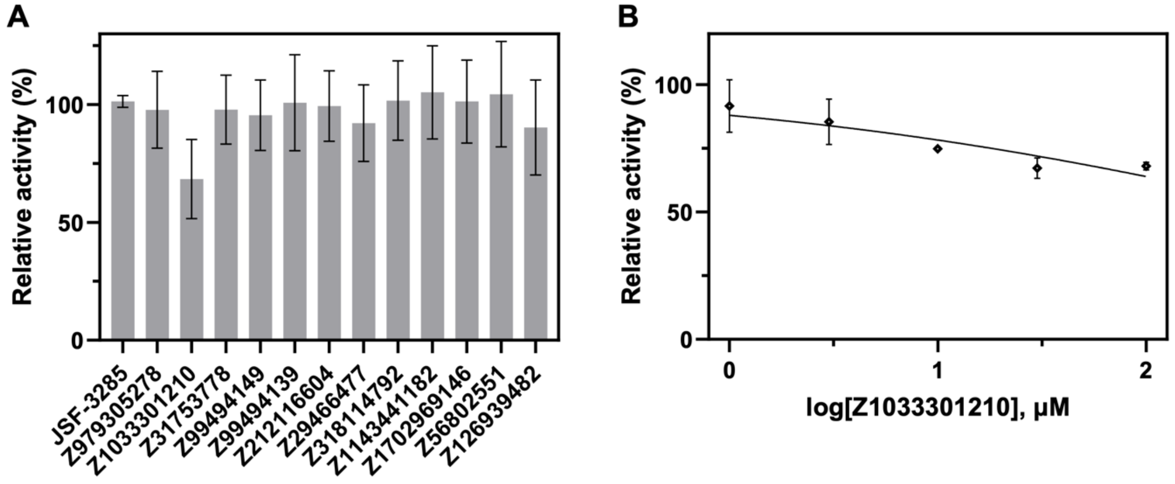

**Figure S2. JSF-5071 mouse po and iv PK profile.** The oral administration route used a solution formulation of 1:9 DMSO:20% Solutol HS15. Each data point represents the mean value ± standard deviation from three mouse experiments.

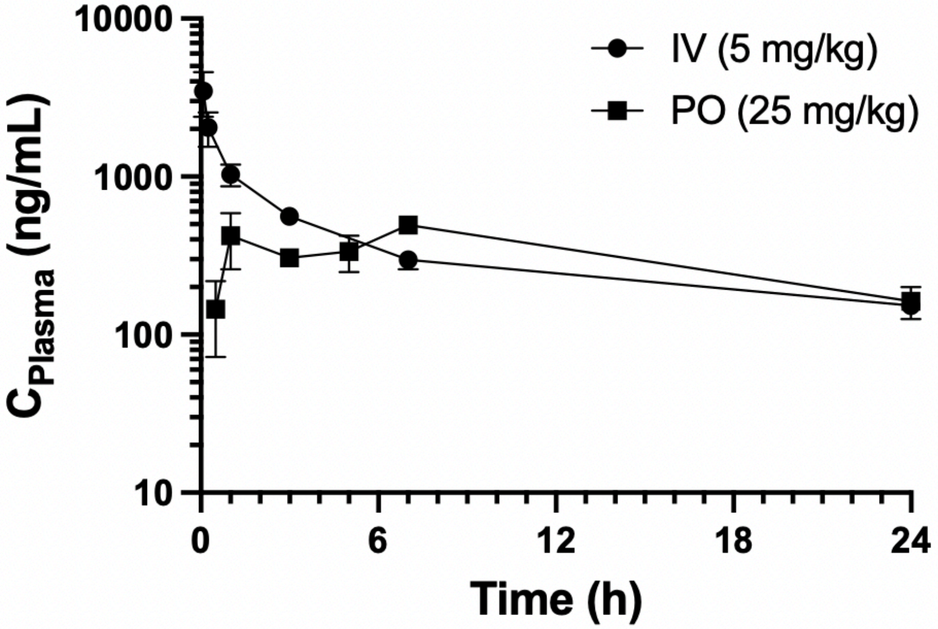

**Figure S3. Dose tolerability mouse PK study for JSF-5071 with data for day 4/4 dosing.** Each data point represents the mean value ± standard deviation from three mouse experiments with 25 mg/kg qd po dosing. The $AUC_{0\text{-}24h}$ was 23,952 h*ng/mL.

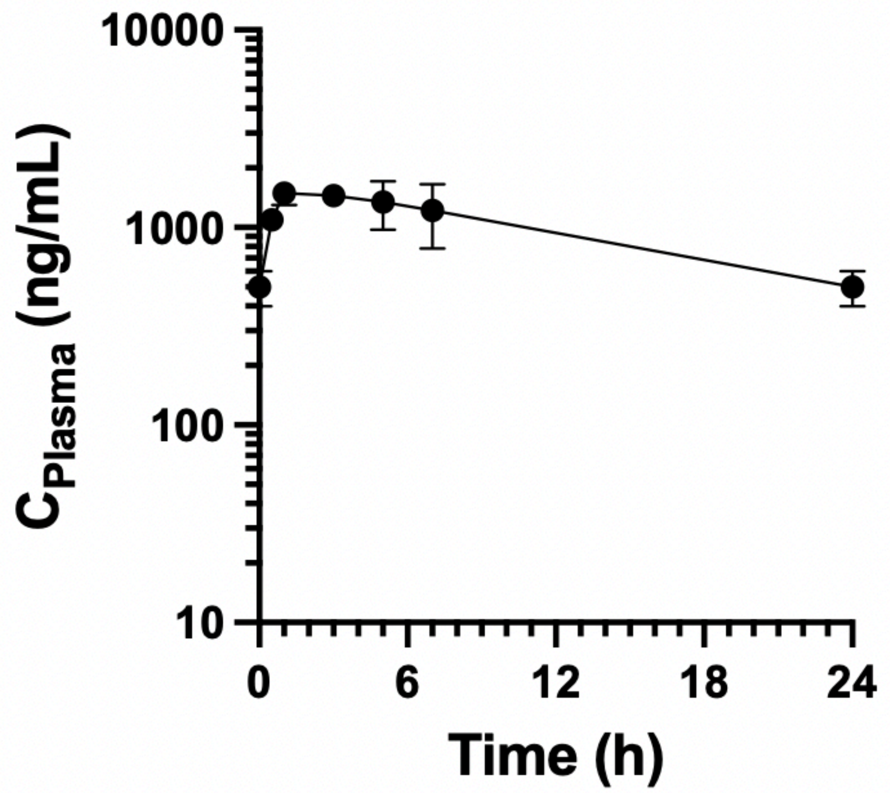

**Figure S4. Evaluation of JSF-5071 in a mouse acute model of *M. tuberculosis* infection with qd dosing.** Bacterial lung burden is shown at 21 d post-infection (dpi). All dosing was administered po bid starting 7 dpi until 18 dpi (12 consecutive d). No-drug controls were administered vehicle only. Each group represents data from five mice. Error bars represent mean ± standard deviation. Ordinary one-way ANOVA with Tukey's *post hoc* multiple comparisons test was used for individual statistical comparisons of all group means. The data were plotted and analyzed using GraphPad Prism 10.2.2. LOD = limit of detection for CFUs.

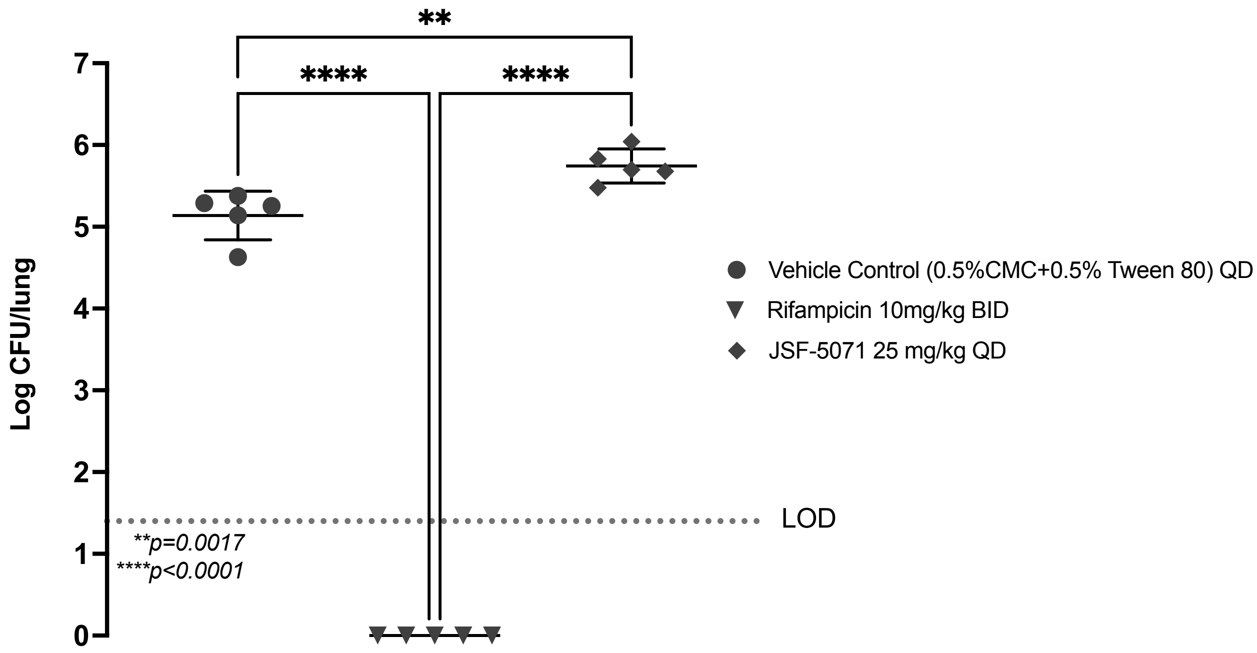


.

**Figure S5. Characterization of isolated mixture of the acyl-AcpM and holo-AcpM species.** (A) Representative total ion chromatogram from LC/MS analysis of Acyl-AcpM mixture used in the KasA functional assay. (B) Distribution of AcpM species across 5 representative preparations.

A

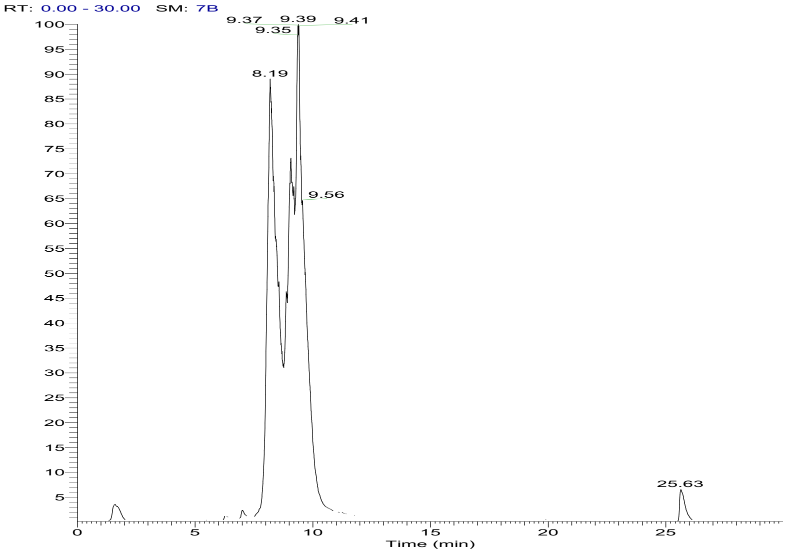


B

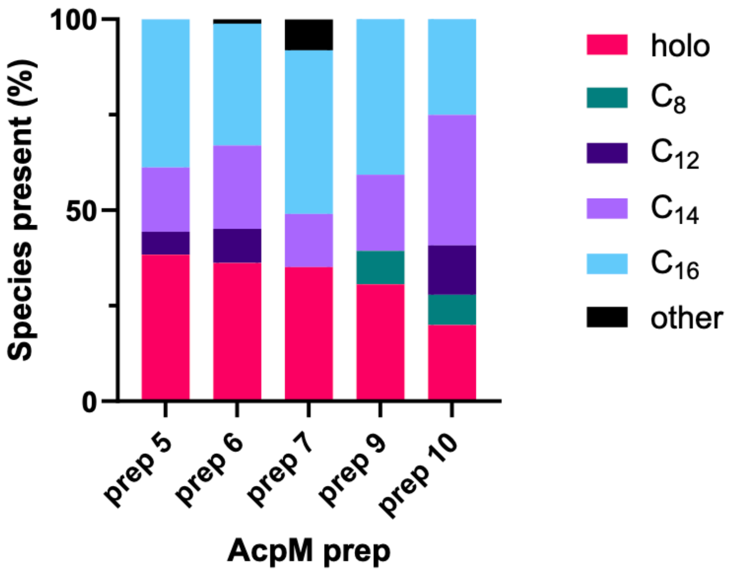

**Table S1. Performance of select models on the experimentally validated 93 compounds, evaluated retrospectively.** An underlined entry indicates the model with the largest value for the pertinent metric. An underlined entry indicates the model with the largest value for the pertinent metric. Models are sorted in descending order of experimental AUROC. The distinguishing details of the named models and their respective training process are provided in Table S2 and the Additional Computational Details: Compound Nomination section below.

| Model | Test AUROC | Experimental AUROC | Experimental Avg. Precision (AP) | Experimental Accuracy |
|---|---|---|---|---|
| | | PK | | |
| PK_RandomForest_1 | 90.3 | 61.9 | 50.9 | 65.2 |
| PK_RandomForest_2 | 90.8 | 54.3 | 46.9 | 57.6 |
| PK_LGBM | 86.4 | 52.6 | 50.3 | 38.0 |
| PK_RandomForest_3 | 93.0 | 51.6 | 46.8 | 58.7 |
| PK_D-MPNN | 85.0 | 49.4 | 40.7 | 53.3 |
| PK_CatBoost | 89.7 | 48.2 | 42.5 | 58.7 |
| PK_RandomForest_4 | 91.1 | 45.7 | 41.2 | 58.7 |
| | | MIC | | |
| MIC_D-MPNN_1 (MLSMR dataset) | 88.6 | 61.9 | 26.9 | 73.9 |
| MIC_RandomForest_1 (MLSMR dataset) | 86.7 | 52.8 | 24.0 | 66.3 |
| MIC_RandomForest_2 (MLSMR dataset) | 83.4 | 51.6 | 21.8 | 67.4 |
| MIC_D-MPNN_2 (MLSMR dataset) | 89.7 | 46.8 | 20.5 | 77.2 |
| MIC_RandomForest_3 (TAACF dataset) | 89.1 | 47.7 | 23.5 | 79.3 |
| MIC_RandomForest_4 (TAACF dataset) | 88.0 | 46.0 | 18.2 | 80.4 |

**Table S2. Feature engineering and pipeline details for select models.**

| Model Name | Dataset | Feature Engineering Pipeline | Optimizer | Objective | Treatment |
|---|---|---|---|---|---|
| PK_RandomForest_1 | PK | P1 | scikit-learn (Random Search) | AUROC | None |
| PK_RandomForest_2 | PK | P2 | Hyperopt (TPE) | AUROC | None |
| PK_RandomForest_3 | PK | P2 | Optuna (TPE) | AUROC | None |
| PK_RandomForest_4 | PK | P2 | scikit-learn (Random Search) | AUROC | None |
| PK_LightGBM | PK | P1 | scikit-learn (Random Search) | AUROC | None |
| PK_CatBoost | PK | P2 | scikit-learn (Random Search) | AUROC | None |
| PK_D-MPNN | PK | N/A | Chemprop | BCE | None |
| MIC_RandomForest_1 | MLSMR | Default | scikit-learn (Random Search) | F1 | Class weight added to loss function |
| MIC_RandomForest_2 | MLSMR | Default | scikit-learn (Random Search) | F1 | None |
| MIC_RandomForest_3 | TAACF | Default | scikit-learn (Random Search) | F1 | Class weight added to loss function |
| MIC_RandomForest_4 | TAACF | Default | scikit-learn (Random Search) | F1 | None |
| MIC_D-MPNN_1 | MLSMR | N/A | Chemprop | BCE | None |
| MIC_D-MPNN_2 | MLSMR | N/A | Chemprop | BCE | None |

**Table S3. KasA functional inhibition data for select compounds.**

| Compound | Reported (% inhibition)* | |
|---|---|---|
| | Average | Standard Deviation |
| JSF-3285 | 92.6 | 3.5 |
| Z979305278 | 31.2 | 9.3 |
| Z1033301210 | 46.7 | 12.5 |
| Z31753778 | 12.0 | 8.8 |
| Z99494149 | 12.0 | 9.7 |
| Z99494139 | 11.5 | 7.0 |
| Z212116604 | 14.7 | 9.6 |
| Z29466477 | 15.3 | 12.8 |
| Z318114792 | 4.72 | 3.00 |
| Z1143441182 | 13.9 | 12.0 |
| Z1702969146 | 27.1 | 8.1 |
| Z56802551 | 15.6 | 11.1 |
| Z126939482 | 21.9 | 13.9 |
| JSF-5071 | 12.7 | 4.8 |

* n = 6 independent trials for all compounds except JSF-5071 (n = 3)

**Table S4. MabA functional inhibition data for select compounds.**

| Compound | Reported % MabA Activity* | |
|---|---|---|
| | Average | Standard Deviation |
| JSF-3285 | 101 | 2.5 |
| Z979305278 | 97.8 | 16.3 |
| Z1033301210 | 68.4 | 16.8 |
| Z31753778 | 97.9 | 14.6 |
| Z99494149 | 95.5 | 15.0 |
| Z99494139 | 101 | 20.4 |
| Z212116604 | 99.4 | 15.0 |
| Z29466477 | 92.1 | 16.2 |
| Z318114792 | 102 | 16.8 |
| Z1143441182 | 105 | 19.7 |
| Z1702969146 | 101 | 17.6 |
| Z56802551 | 104 | 22.4 |
| Z126939482 | 90.3 | 20.2 |

* n = 6 independent trials

**Table S5. Mouse PK parameters for JSF-5071 with a formulation of 5% DMA/95%(4% Cremophor) for iv dosing.** The dosing levels for iv and po administration were 5 and 25 mg/kg, respectively.

| Half-Life (h) | $V_d$ (L/kg) | Oral Bioavailability (%) |
|---|---|---|
| 9.72 | 6.34 | 18.5[a], 10.7[b] |

[a]po dosing was with a solution formulation of 1:9 DMSO:20% Solutol HS15
[b]po dosing was with a suspension formulation of 5% DMA/95% CMC/Tween

**Materials and Methods**

**<u>Additional Computational Details</u>**

***Feature Engineering* Pipelines** – We tested and compared several feature transformation pipelines in addition to hyperparameter optimization algorithms. The first feature transformation pipeline 1 ("P1") involves the following steps: 1) Multicollinearity(ρ, 0.9) → Variance Thresholding(0.2) → Normalization. The second pipeline ("P2") involves: Multicollinearity(ρ, 0.95) → Variance Thresholding(0.1) → Importance-Score → Normalization. Additional details are provided in the main text Materials and Methods. Model hyperparameters were optimized using scikit-learn (for random search) and Hyperopt and Optuna (for Bayesian search using the TPE algorithm).

*Training* – Models were trained using a weighted binary cross-entropy loss function. For the PK dataset, the sample weights were not varied when calculating the loss. For the MIC datasets, weights for negative samples were fixed to 1, and weights for positive samples were determined using the hyperparameter optimization procedure. All neural models were built using PyTorch and PyTorch Lightning and trained using a SGD optimizer. The momentum $\mu$ was set to 0.9 and no weight decay or dampening was used ($\lambda = 0$ and $\tau = 0$). The batch size was set to 2048 for the MIC datasets and 100 for the PK dataset. The training/fine-tuning process employed an early stopping mechanism with a tolerance of 7 iterations. The learning rate was determined through hyperparameter optimization. The tokenizer sequence length was fixed to 128, i.e., sequences shorter than 128 tokens were padded to this length, and sequences longer than 128 tokens were truncated.

*Hyperparameter Optimization* – Hyperparameter optimization was performed with the Tree-Structured Parzen Estimator and Random Search optimization methods available in the Python libraries Hyperopt, Optuna, and scikit-learn. We tested various objective functions including the area under the Receiver Operating Characteristic curve (AUROC), Youden's J, and maximum F1-score. The exact settings used in the models for compound nomination are listed in Table S2. Architectural hyperparameters common for the ChemBERTa-v2 model include i. the number of layers in the fully connected classification head and ii. the learning rate. The embedding dimension was fixed at 384 and the sizes of the fully connected layers were determined as follows: given a starting dimension and the number of layers, we cascaded the layer sizes in powers of 2. We used the Optuna library and validation loss as the objective function. We ran a study of 20 trials for the

PK dataset, 10 trials for the MIC-TAACF dataset, and 10 trials for the MIC-MLSMR dataset with five warmup trials each to arrive at an initial set of architecture and training hyperparameters. These hyperparameters were then tuned manually by varying each hyperparameter independently. In each trial, the model instance was trained for 50 epochs with early stopping. Hyperparameter search for the D-MPNN models was performed with the Chemprop package, which utilizes Hyperopt. The searched hyperparameters included the number of hidden layers in the readout phase, hidden size of message vectors, dropout rate, and depth (number of message-passing layers). The optimization was executed for 10, 10, and 20 iterations for the PK, MIC-TAACF, and MIC-MLSMR datasets, respectively, with BCE loss as the objective function. The batch size was fixed to 32 samples for the MIC datasets and 16 samples for the PK dataset. Each trial was run for 200 epochs for the MIC-TAACF dataset and 100 epochs for the other two datasets.

*Compound Nomination* – For PK compound scoring, we chose a combination of Random Forest, LightGBM, CatBoost, and D-MPNN models based on their test performance and checked for overfitting by comparing it with their training set performance. The details of the selected models are presented in Table S2.

Multi-objective Optimization – With multiple surrogate model scores available, downselection from the set of ($N$ = 956) compounds selected in the docking step was treated as a multi-objective optimization problem. For each model $m$ predicting $g \in \{\mathrm{PK}, \mathrm{MIC}\}$ and each compound $i$, let $y_m^g(i) \in \{0,1\}$ represent the binary model prediction and $r_m^g(i) \in \{1, \dots, N\}$ represent the rank given to the molecule by the model. Let there be a total of $n_{\mathrm{PK}}$ PK models and $n_{\mathrm{MIC}}$ MIC models. The task is then to select top $k$ compounds that are predicted to fulfill MIC and PK criteria according to multiple model instances. To achieve this goal, we defined three multiparameter scores and selected compounds with each.

The first multi-parameter score $PS_{\mathrm{all}}$ represents the number of all models (PK and MIC) that predict a compound $i$ to be active:

$$PS_{\mathrm{all}} = \sum_{g \in \{\mathrm{MIC}, \mathrm{PK}\}} \sum_{m=1}^{n_g} y_m^g(i).$$

We selected 31 compounds using this score, referred to as the “Majority Voting by All the Models” selection strategy.

The second multi-parameter score $NSR_{\text{all}}$ is the negative sum of the ranks of all models. This score will be maximized for compounds which are top-ranking according to all PK and MIC models:

$$NSR_{\text{all}} = -\sum_{g\in\{\text{MIC,PK}\}} \sum_{m=1}^{n_g} \text{rank}_m^g(i).$$

The 31 compounds that optimized this score were selected as the "Negative Sum of Ranks of All the Models" selection.

The third multi-parameter score $PS_{\text{MIC}}$ represents the number of MIC models that predict a compound $i$ to be active:

$$PS_{\text{MIC}} = \sum_{m=1}^{n_{\text{MIC}}} y_m^{\text{MIC}}(i).$$

In the main text, we refer to this as "Majority Voting by the MIC Models". We selected 10 compounds that optimized this multiparameter score.

The sets of 31, 31, and 10 compounds selected using each score resulted in an initial set of 47 unique molecules.

*Diversity-Based Selection* – The top 956 compounds were clustered into 20 groups by structural similarity with k-means clustering using ECFP4 fingerprints as molecular features. The initial set of 47 selected compounds only covered 11 of the 20 clusters. To increase the diversity in the downselected set, we also chose the top 3 compounds sorted by $NSR_{\text{all}}$ from the remaining 9 clusters, resulting in the addition of 27 compounds to the initial set. To further maximize the chemical space coverage, we selected one or two compounds from each cluster providing an additional 22 compounds, considering the following factors:

• If the conformer appeared reasonable upon inspection, emphasis was placed on occupying the hydrophobic channel where the JSF-3285 4-fluorobutyl group was located;

• Since Glu-199 plays a crucial role in hydrogen bonding with JSF-3285, preference was given to compounds proposed to form a hydrogen bond with Glu-199;

• Proposed pi-pi face-to-face interactions with Phe-239 were deemed favorable;

• Proposed hydrogen bonding with Gly-200 and/or Ala-119;

• When multiple compounds met all the above criteria, lower molecular weight compounds were prioritized due to their potential for greater ligand efficiency and in consideration of their future optimization where molecular weight could be increased.

Overall, the three selected sets yielded 47 (model-based), 27 (model + diversity-based) and 22 (inspection + diversity-based) compounds respectively, with a total of 96 compounds. Three compounds (Z56760621, Z56822324, Z56799038) of these 96 were not sourced due to issues with the supplier regarding excessive cost and/or delivery time.

### Additional Compound Characterization Data

Given the downstream *in vivo* assessment of JSF-5071, 5077, 5088, and 5089, we have included an HPLC trace for each compound which demonstrated >99%, >99%, >99%, and >99% purity, respectively at 250 nm. The remainder of their characterization data may be found in the Experimental Section of the main text.

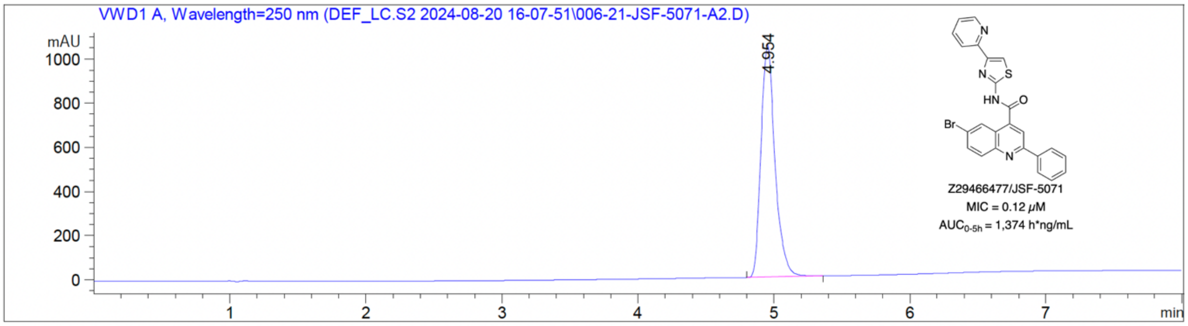


```
Signal 2: VWD1 A, Wavelength=250 nm

Peak RetTime Type  Width     Area       Height     Area
  #   [min]        [min]   [mAU*s]      [mAU]        %
----|-------|----|-------|----------|----------|--------|
   1   4.954 BB    0.1103 7525.66943 1053.83838 100.0000

Totals :                  7525.66943 1053.83838
```

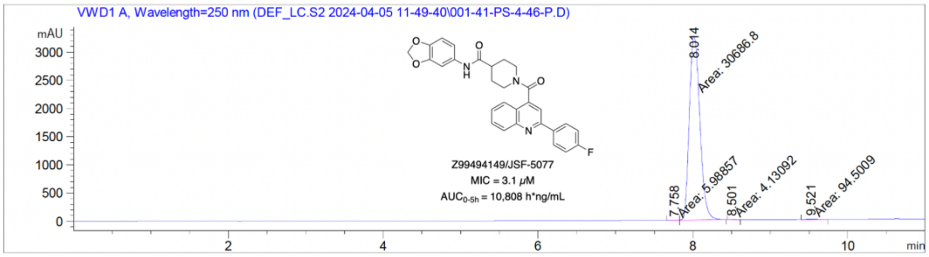


Signal 2: VWD1 A, Wavelength=250 nm

| Peak # | RetTime [min] | Type | Width [min] | Area [mAU*s] | Height [mAU] | Area % |
|---|---|---|---|---|---|---|
| 1 | 7.758 | MM | 0.1086 | 5.98857 | 9.18806e-1 | 0.0194 |
| 2 | 8.014 | MM | 0.1568 | 3.06868e4 | 3261.32446 | 99.6602 |
| 3 | 8.501 | MM | 0.0968 | 4.13092 | 7.11429e-1 | 0.0134 |
| 4 | 9.521 | MM | 0.1836 | 94.50092 | 8.58070 | 0.3069 |

Totals : 3.07914e4 3271.53540

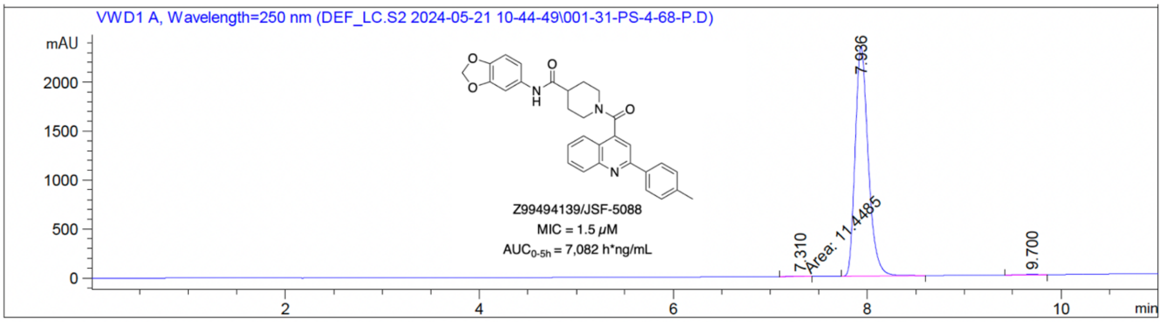


Signal 2: VWD1 A, Wavelength=250 nm

| Peak # | RetTime [min] | Type | Width [min] | Area [mAU*s] | Height [mAU] | Area % |
|---|---|---|---|---|---|---|
| 1 | 7.310 | MM | 0.2333 | 11.44854 | 8.17908e-1 | 0.0525 |
| 2 | 7.936 | BB | 0.1423 | 2.17460e4 | 2342.69873 | 99.7353 |
| 3 | 9.700 | BB | 0.2018 | 46.26925 | 3.51025 | 0.2122 |

Totals : 2.18037e4 2347.02689

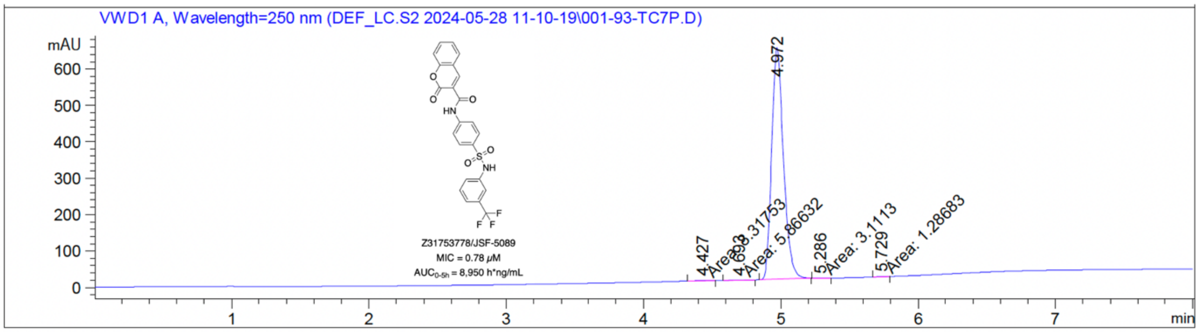


Signal 2: VWD1 A, Wavelength=250 nm

| Peak # | RetTime [min] | Type | Width [min] | Area [mAU*s] | Height [mAU] | Area % |
|---|---|---|---|---|---|---|
| 1 | 4.427 | MM | 0.0953 | 8.31753 | 1.45440 | 0.2179 |
| 2 | 4.693 | MM | 0.1526 | 5.86632 | 6.40901e-1 | 0.1537 |
| 3 | 4.972 | BB | 0.0908 | 3798.56299 | 635.19482 | 99.5132 |
| 4 | 5.286 | MM | 0.0717 | 3.11130 | 7.23208e-1 | 0.0815 |
| 5 | 5.729 | MM | 0.0664 | 1.28683 | 3.23175e-1 | 0.0337 |

Totals : 3817.14498 638.33651
